# Synchrotron Nano-FTIR Reveals Carbohydrate-Dependent Protein Conformational Changes at Bacterium–Nanoparticle Interfaces

**DOI:** 10.64898/2026.08.18.745527

**Authors:** Clara Lana Bispo Fidelis, Aline Orvalho Pereira, Renata Santos Rabelo, Lindomar José Calumby Albuquerque, Luelc Souza da Costa, Ohanna Maria Menezes Madeiro da Costa, Jefferson Bettini, Raul de Oliveira Freitas, Mateus Borba Cardoso

## Abstract

Antimicrobial resistance motivates the development of approaches capable of probing nanoparticle–bacterium interactions with nanoscale sensitivity. Here, synchrotron infrared nano-spectroscopy (SINS) is applied to investigate interactions between carbohydrate-coated silica nanoparticles and the Gram-negative model bacterium *Escherichia coli* at the single-cell level. Silica nanoparticles (SiO_2_) were coated with mannose, maltose, or trehalose to evaluate how surface carbohydrate chemistry influences their interactions with the bacterial envelope. Correlative electron microscopy revealed pronounced association of carbohydrate-SiO_2_ with the bacterial envelope, with features consistent with localization within the periplasmic region, whereas bare-SiO_2_ showed no detectable association. SINS measurements acquired directly on bacterial cells and at bacterium–nanoparticle interfaces revealed distinct, carbohydrate-dependent spectral signatures. Quantitative analysis of the amide I band used the Iα/Iβ ratio, which describes the relative contributions of α-helical and β-sheet protein secondary-structure components, together with interface-dependent band-position analysis to characterize local spectral perturbations. Carbohydrate-SiO_2_ produced systematic changes in the Iα/Iβ ratio, including at locations where nanoparticles were not directly observed, indicating that their effects extend beyond the sites of nanoparticle association. Comparison of measurements acquired on bacterial surfaces and at bacterium–nanoparticle interfaces further revealed that carbohydrate chemistry modulates both the magnitude and spatial extent of these spectral perturbations. Trehalose-SiO_2_ produced the largest interface-dependent amide I band shifts and a spectral component consistent with random-coil structures. Overall, these results demonstrate that carbohydrate surface chemistry modulates nanoscale protein conformational perturbations at the nano–bio interface and highlight SINS as a powerful approach for resolving chemically localized molecular responses at single-cell interfaces.

## INTRODUCTION

Antimicrobial resistance represents an urgent global health crisis with profound societal and economic impacts ^1–4^. In response, nanobiotechnology has emerged as a versatile platform for extending drug half-life, improving delivery, enhancing microbial detection, and promoting direct interactions with bacterial envelopes ^5–8^. Despite these advances, experimental tools capable of resolving with molecular specificity how nanoparticle interactions modify the local chemical environment of bacterial envelopes at the nanoscale remain limited. ^9,10^.

Gram-negative (GN) bacteria pose a particular challenge for nano–bio investigations due to their outer membrane (OM), a highly organized and selective barrier composed of lipopolysaccharides and membrane proteins that regulate molecular transport ^11,12^. This structural complexity limits nanoparticle access and complicates direct interaction, especially in the absence of clear anchoring sites at the cell surface. While several targeting strategies have been explored—such as antibiotics, peptides, and membrane-derived vesicles—most successful examples have focused on Gram-positive bacteria ^8,13–15^.

Recent studies have highlighted the role of glycan-mediated interactions between the GN OM and host tissues, revealing that carbohydrate recognition can facilitate bacterial adhesion and transport processes ^16^. This insight has opened new opportunities in nanomedicine, where glycan-functionalized nanoparticles may exploit natural recognition pathways to promote controlled interactions with GN bacteria ^17,18^. However, despite growing interest, direct experimental evidence linking glycan-mediated nanoparticle binding to nanoscale perturbations of the bacterial OM remains scarce.

A major obstacle to progress lies in the limitations of existing characterization techniques. High-resolution electron microscopy provides detailed morphological information but lacks chemical specificity, whereas conventional infrared (IR) spectroscopy captures biochemical fingerprints only at the bulk or micrometer scale ^19–21^. Although synchrotron-based micro-FTIR improves sensitivity ^22–26^, it remains insufficient to resolve nanoscale heterogeneities at the bacterium–nanoparticle interface. Beyond spatial resolution limitations, an additional challenge arises from the extreme sensitivity of biomolecular functional groups to their local chemical environment. At nanoparticle–membrane interfaces, nanoscale confinement can substantially modify vibrational behavior, producing spectral signatures that do not emerge in bulk or ensemble measurements and remain inaccessible to diffraction limited approaches. Resolving these interfacial chemical identities therefore requires analytical probes capable of operating directly at the spatial scale where such confinement driven effects originate.

Infrared nano-spectroscopy overcomes these limitations by combining atomic force microscopy with localized infrared excitation, enabling chemical analysis with spatial resolution down to tens of nanometers ^27–30^. In particular, synchrotron-based nano-FTIR provides the sensitivity required to probe single bacterial cells and their interfaces with nanomaterials, enabling direct access to localized vibrational signatures that arise specifically from confined nano–bio interactions ^31–33^. While early work, including our previous study, demonstrated the feasibility of applying nano-IR techniques to nanoparticle–cell systems ^17^, a systematic investigation of how nanoparticle surface chemistry modulates nanoscale-confined spectral perturbations at bacterial membranes has not yet been established.

Here, we investigate how carbohydrate coating of silica nanoparticles with mannose, maltose, or trehalose influences protein conformational changes at the nano–bio interface using synchrotron infrared nano-spectroscopy (SINS). By correlating electron microscopy with SINS measurements acquired directly on bacterial cells and at bacterium–nanoparticle interfaces, we demonstrate that nanoparticle surface chemistry modulates the magnitude and spatial extent of protein conformational remodeling. These results highlight the unique capability of SINS to resolve chemically confined molecular responses within the nanometric interaction volume of the near field, providing access to interface-specific molecular information that remains inaccessible to ensemble-averaged and diffraction-limited spectroscopic techniques. ^27–30^. This work establishes SINS as a powerful platform for investigating how nanoparticle surface chemistry shapes molecular processes at biologically relevant nano–bio interfaces.

## RESULTS AND DISCUSSION

In this work, silica nanoparticles (SiO_2_) were obtained by conventional one-pot Stöber synthesis ^34^. Bare-SiO_2_ was incubated with carbohydrates mannose, maltose and trehalose to obtain surface-modified nanoparticles with each individual sugar (man-SiO_2_, mal-SiO_2_ and tre-SiO_2_, respectively). After 1 hour of incubation, the suspension was centrifuged, and the supernatant was discarded to remove the non-adsorbed carbohydrates. After 2 washes, no free carbohydrate was detected in the supernatant by the consolidated phenol-sulfuric acid method ^35^. Data on free carbohydrate removal are shown in Fig. S1 (Supporting Information). Bare-SiO_2_ and freshly prepared carbohydrate coated nanoparticles (carb-SiO_2_) underwent physical-chemical characterization. Scanning Transmission Electron Microscopy (STEM) was employed to investigate the diameter and morphology of the nanoparticles (NPs) (Fig. S2, Supporting Information). STEM images revealed uniformly sized spherical particles with an average diameter of approximately 70 nm, independent of sample composition. Colloidal stability was assessed by Dynamic Light Scattering (DLS) in water, PBS, DMEM, and LB broth (Fig. S3–S5, Supporting Information). Nanoparticles remained stable in water, PBS, and DMEM over the evaluated time window, whereas aggregation was observed in LB broth, a highly complex medium. Although subtle aggregation occurs under these conditions, bacterial incubations were short-term and followed by washing steps prior to imaging and spectroscopy, ensuring that the observed nanoparticle–bacterium interactions reflect surface chemistry rather than bulk aggregation effects.

Energy Filtered Transmission Electron Microscopy (EFTEM), a technique that combines the high resolution of TEM with electron energy loss spectroscopy (EELS), enabled the chemical samples’ characterization by detecting inelastic electron scattering ^36^. Consequently, elemental map images can be generated by detecting energy loss corresponding to inner-shell ionization ^37^. Fig. 1 shows EFTEM images of bare-SiO_2_ and carb-SiO_2_, featuring elemental maps for Si represented in red and C labeled in green. The noticeable green signal originated primarily from the holey carbon TEM grids. Additionally, a green signal was detected in all NPs, indicating organic content. However, carb-SiO_2_ exhibited a distinct carbon corona surrounding the NPs irrespective of the carbohydrate-coated sample. These observations confirmed the presence of an organic corona consistent with carbohydrate, thereby validating our adsorption method. These images could discriminate between bare and carbohydrate-coated particles, reinforcing our DLS results.

**Figure 1.**
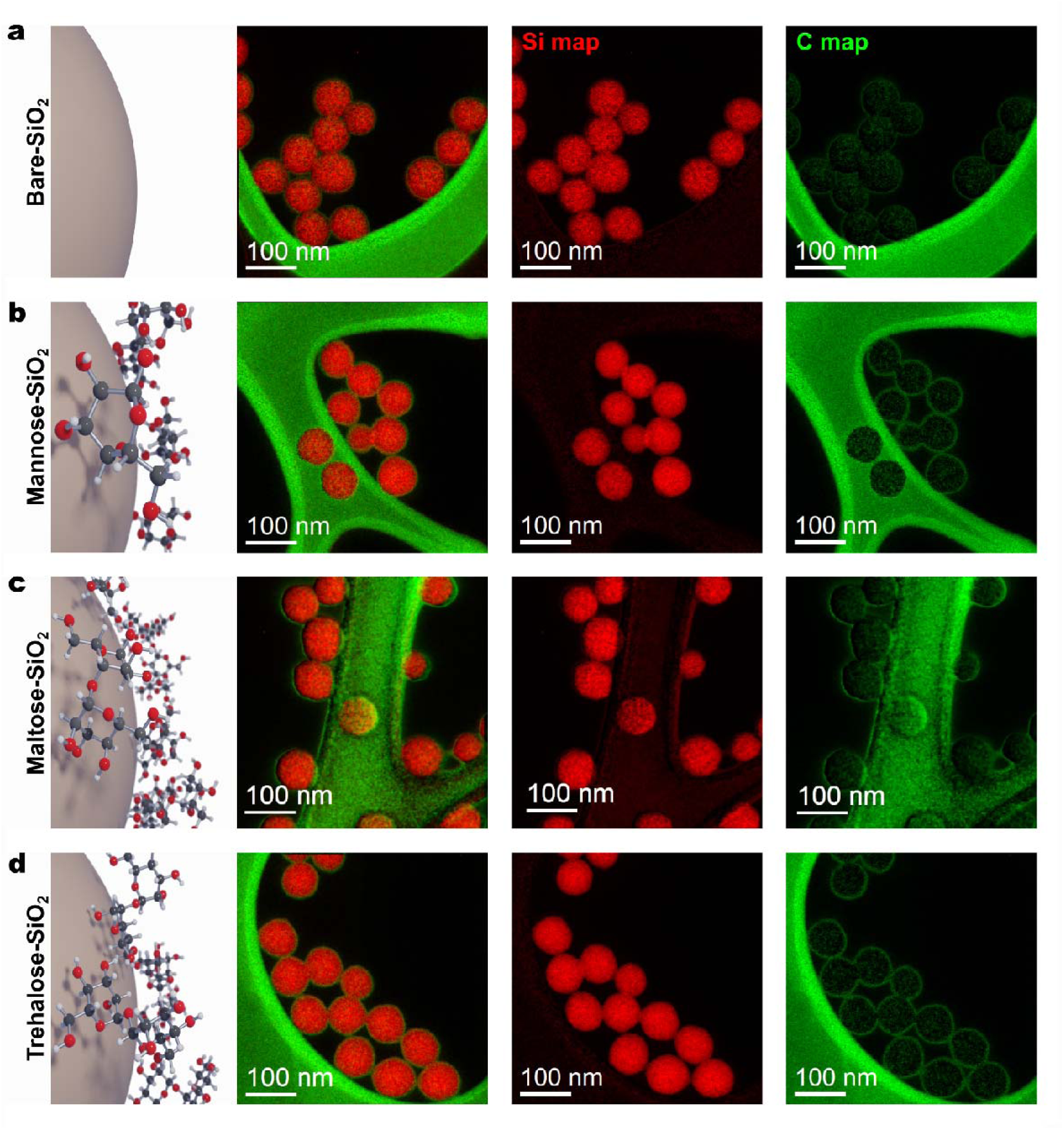
EFTEM elemental maps of NPs. Distribution of Si (red) and C (green) for (a) bare-SiO_2_, (b) man-SiO_2_, (c) mal-SiO_2_, and (d) tre-SiO_2_.

Biological assays were performed to evaluate bacterial growth perturbation in *E. coli* and cell viability in mammalian cells (NIH-3T3 fibroblasts). The bacterial growth assay, presented in Fig. S6, involved cultivating *E. coli* with distinct nanoparticle concentrations for 24 hours, followed by optical density measurements to estimate cell growth. Bare-SiO_2_ did not affect bacterial growth across the tested concentrations, whereas carb-SiO_2_ induced a reduction in *E. coli* growth. Statistical analysis was conducted using two-way ANOVA to compare bacterial growth in the presence of bare and carbohydrate-coated nanoparticles, as detailed in Table S2.

A carbohydrate-dependent trend was observed, with trehalose-, mannose-, and maltose-coated nanoparticles producing different degrees of reduction in bacterial growth, with trehalose showing the strongest effect. These results indicate that nanoparticle surface coating modulates bacterial response upon exposure and are consistent with preferential interactions between carbohydrate-coated nanoparticles and the bacterial envelope. While specific molecular pathways are not resolved here, the observed trends align with reported affinities between carbohydrates and bacterial surface components described in the literature ^18,38–43^ .

In parallel, nanoparticle effects on mammalian cells were assessed using NIH-3T3 fibroblasts. Cell viability assays (Fig. S7 and Table S3, Supporting Information) showed that all nanoparticle formulations were well tolerated up to 125 μg mL^-1^, with reduced viability observed only at higher concentrations. Carbohydrate-coated nanoparticles exhibited slightly higher cell viability compared to bare-SiO_2_ under these conditions, consistent with previous reports of carbohydrate-mediated modulation of nanoparticle–cell interactions ^44–47^. These assays provide a comparative reference between bacterial and mammalian responses under identical exposure conditions.

Scanning electron microscopy (SEM) was employed to evaluate nanoparticle association with the bacterial OM. *E. coli* was incubated separately with the four types of nanoparticles (bare, man-, mal-, and tre-SiO_2_) for 15 minutes at a concentration of 100 μg mL^-1^ per 10□ CFU mL^-1^ of cells. The suspensions were washed to remove residual culture medium and salts and deposited onto glass substrates pre-treated with poly-L-lysine. The cells were subsequently fixed and dried by CO□-critical-point drying. SEM images show that bare-SiO_2_ predominantly forms micrometric aggregates with no detectable association to the bacterial surface (Fig. 2a). In contrast, carb-SiO_2_ are consistently observed in close proximity to the bacterial envelope (Fig. 2b–g), remaining associated after extensive washing and drying steps. Representative images from different substrate regions are shown, and the observations were reproducible across independent sample preparations.

**Figure 2.**
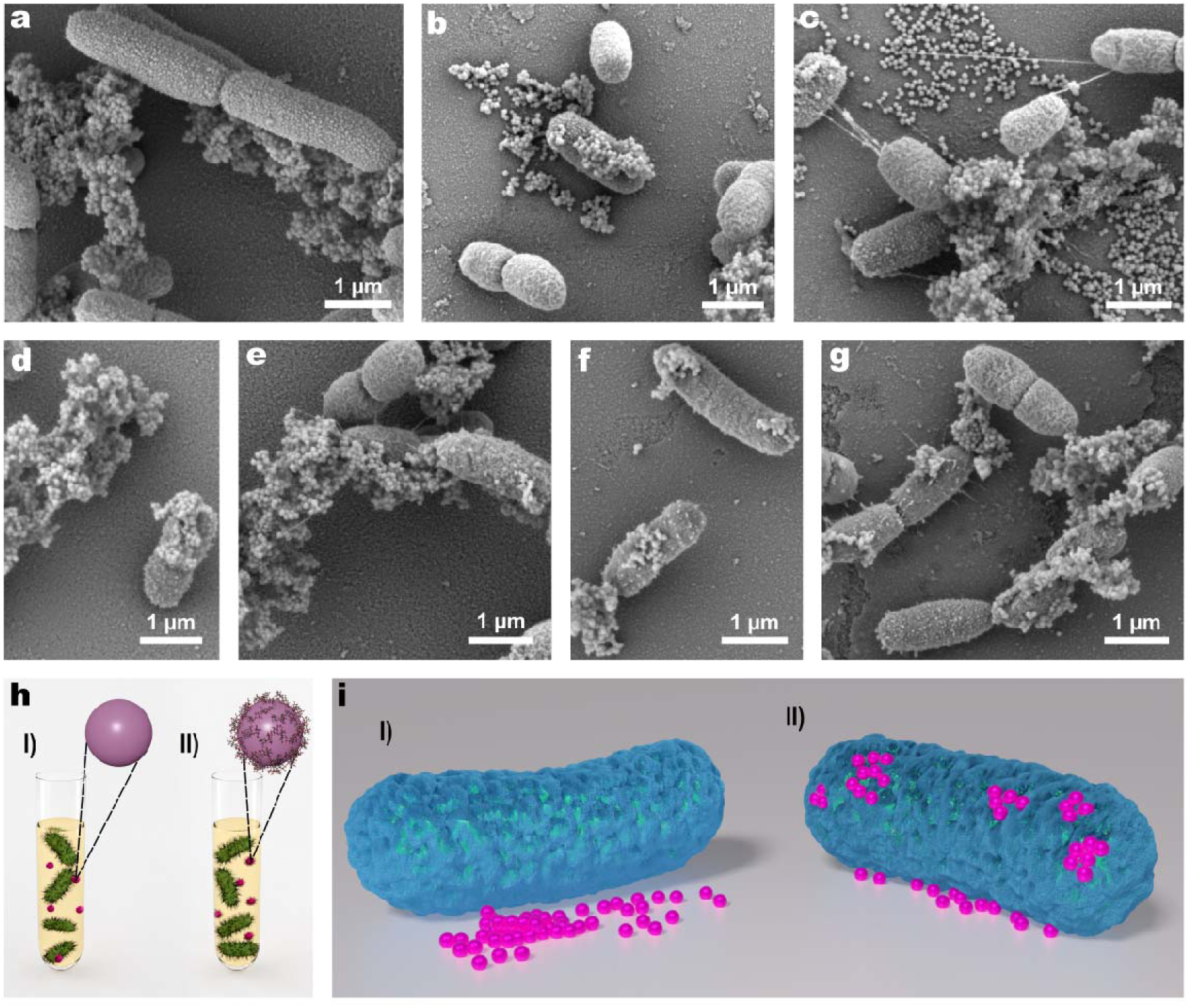
**SEM analysis of *E. coli*–nanoparticle interactions. (**a–g) SEM images of *E. coli* incubated with NPs. (a) bare-SiO_2_, showing no evident bacterium–nanoparticle interaction. (b,c) man-SiO_2_; (d,e) mal-SiO_2_; and (f,g) tre-SiO_2_, showing structures consistent with nanoparticle association at the bacterial surface. Scale bars are indicated in the images. (h) Schematic representation of the two incubation conditions used in this study: (I) *E. coli* incubated with bare-SiO_2_; and (II) *E. coli* incubated with carb-SiO_2_. (i) Schematic illustration of the corresponding bacterial configurations: (I) absence of bacterium–nanoparticle interaction; and (II) nanoparticle association at the bacterial surface.

Fig. 2h schematically summarizes the two incubation conditions, while Fig. 2i illustrates the corresponding outcomes. Whereas bare-SiO_2_ shows no detectable surface association (Fig. 2i–I), carbohydrate-coated nanoparticles display pronounced surface localization at the GN bacterial envelope (Fig. 2i–II), consistent with a carbohydrate- dependent interaction.

Transmission electron microscopy (TEM) was employed to investigate nanoparticle–cell associations at higher spatial resolution. TEM is particularly suited for evaluating bacterial ultrastructure, i.e., the nanoscale organization of cellular components ^48,49^. Samples were prepared using the same incubation conditions as for SEM, followed by osmium tetroxide staining, resin embedding, and sectioning into 70–100 nm slices using an ultramicrotome. Representative thin sections are shown in Fig. 3.

**Figure 3.**
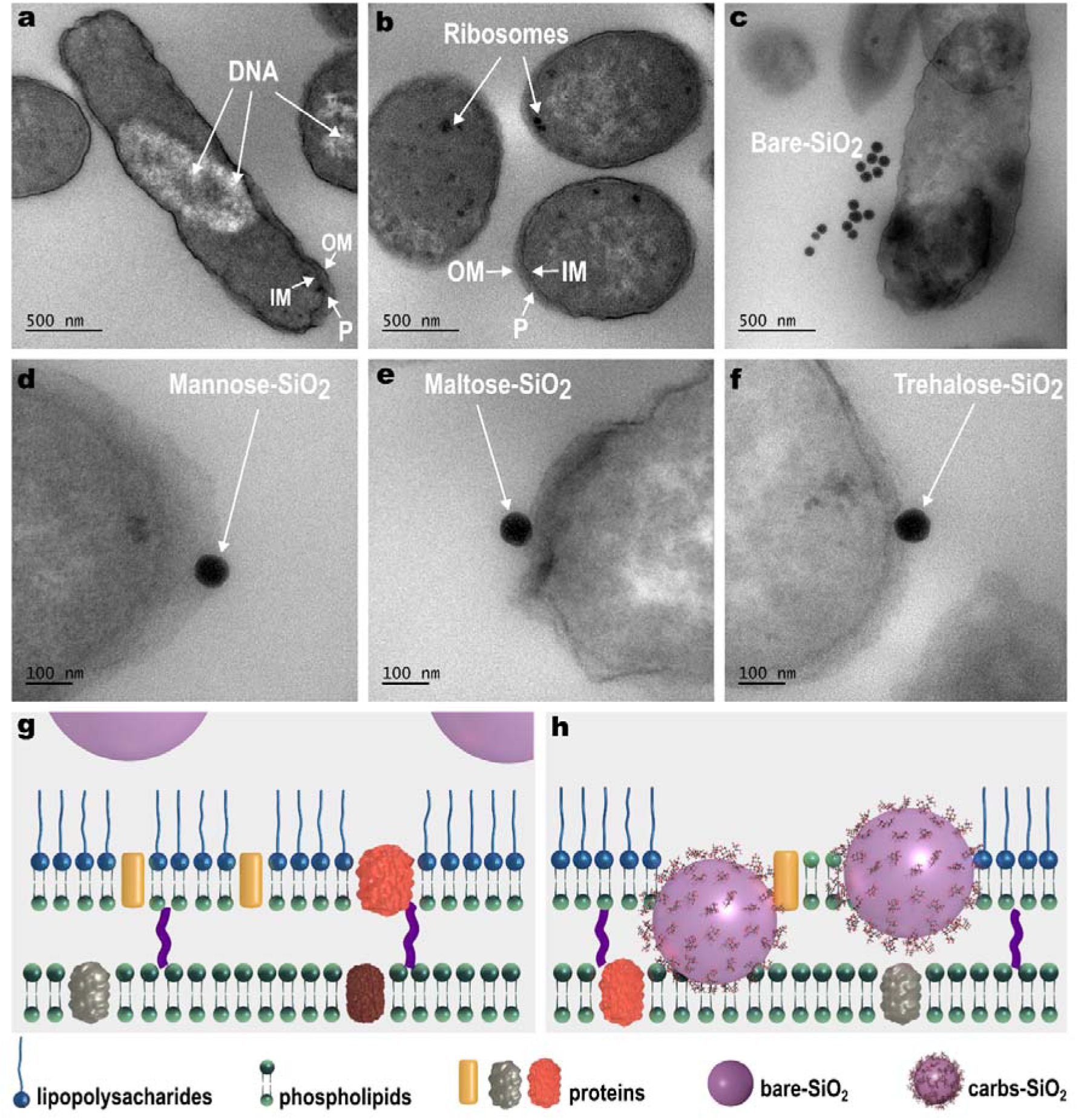
TEM analysis of *E. coli* incubated with NPs. (a,b) Control *E. coli* cells in the absence of NPs. White arrows indicate cellular features, including DNA, ribosomes, and membrane structures (OM, outer membrane; IM, inner membrane; P, periplasm). (c) *E. coli* incubated with bare-SiO_2_. High-magnification TEM images of *E. coli* incubated with (d) man-SiO_2_, (e) mal-SiO_2_, and (f) tre-SiO_2_. (g,h) Schematic representations of the *E. coli* outer membrane following incubation with NPs: (g) preservation of membrane organization in cells incubated with bare-SiO_2_ and (h) membrane alterations associated with incubation with carb-SiO_2_.

Control bacteria incubated without nanoparticles exhibited a typical rod-shaped morphology (Fig. 3a). Electron-light regions corresponding to genetic material and electron-dense regions corresponding to ribosomes are visible within the cytoplasm. Due to variations in sectioning orientation, the OM, peptidoglycan layer, and inner membrane are not always clearly resolved along the entire cell perimeter. In some sections, a wavy OM and a discernible periplasmic space above the inner membrane can be observed (Fig. 3a-b) ^50^. The periplasm corresponds to the compartment between the outer and inner membranes of GN bacteria and mediates transport between the cell interior and exterior ^49–51^.

TEM images of *E. coli* incubated with bare-SiO_2_ show no detectable nanoparticle association with the bacterial envelope, consistent with SEM observations (Fig. 3c). In contrast, high-magnification images of carb-SiO_2_ (Fig. 3d–f) reveal individual nanoparticles localized within the periplasmic region in thin sections. Fig. 3g and Fig. 3h illustrate two distinct scenarios: one in which bare-SiO_2_ does not induce detectable alterations in bacterial ultrastructure, and another in which carbohydrate-coated nanoparticles are associated with localized changes in envelope organization. These observations indicate that carbohydrate coating promotes nanoparticle association with the GN envelope. While the images are consistent with translocation across the OM, they do not resolve the underlying molecular mechanisms. Taken together, SEM and TEM results show that carbohydrate-coated nanoparticles associate with the GN bacterial envelope and localize within the periplasmic region. To further examine nanoparticle–bacterium interactions beyond morphology, SINS was applied.

Further SINS experiments on nanoparticle–bacterium interactions were conducted at the Imbuia beamline at SIRIUS, the fourth-generation synchrotron source in Campinas, Brazil ^32^. SINS combines atomic force microscopy with localized infrared excitation, enabling correlated topographic and infrared measurements with spatial resolution on the order of 25 nm ^31,33^. This capability allows interface-resolved interrogation of individual bacterial cells and nanoparticle-associated regions. In contrast to electron microscopy, SINS provides access to nanoscale-confined spectral perturbations induced by nanoparticle proximity.

*E. coli* was incubated with different nanoparticles, following a sample preparation protocol similar to that used for electron microscopy, and then deposited onto Au-Si substrates. However, for SINS analysis, two crucial considerations must be observed. First, PBS must be avoided during the bacterial washing steps, as it can introduce artifacts into the IR signal. Second, cells must not be chemically fixed, as fixation involves cross-linking between cellular proteins and carbohydrates ^52^, which could lead to inaccurate IR data.

Fig. 4a schematically illustrates the SINS experiment at the Imbuia beamline, in which AFM topography and infrared near-field signals are simultaneously collected under synchrotron infrared illumination of the AFM tip. The incident radiation is concentrated at the tip apex through an antenna effect, enabling nanoscale-resolved spectroscopic measurements ^53^. The resulting near-field interaction generates a scattered signal that encodes the material’s local infrared optical response, enabling nanoscale spectroscopic probing of molecular vibrational features ^31,33,53,54^.

**Figure 4.**
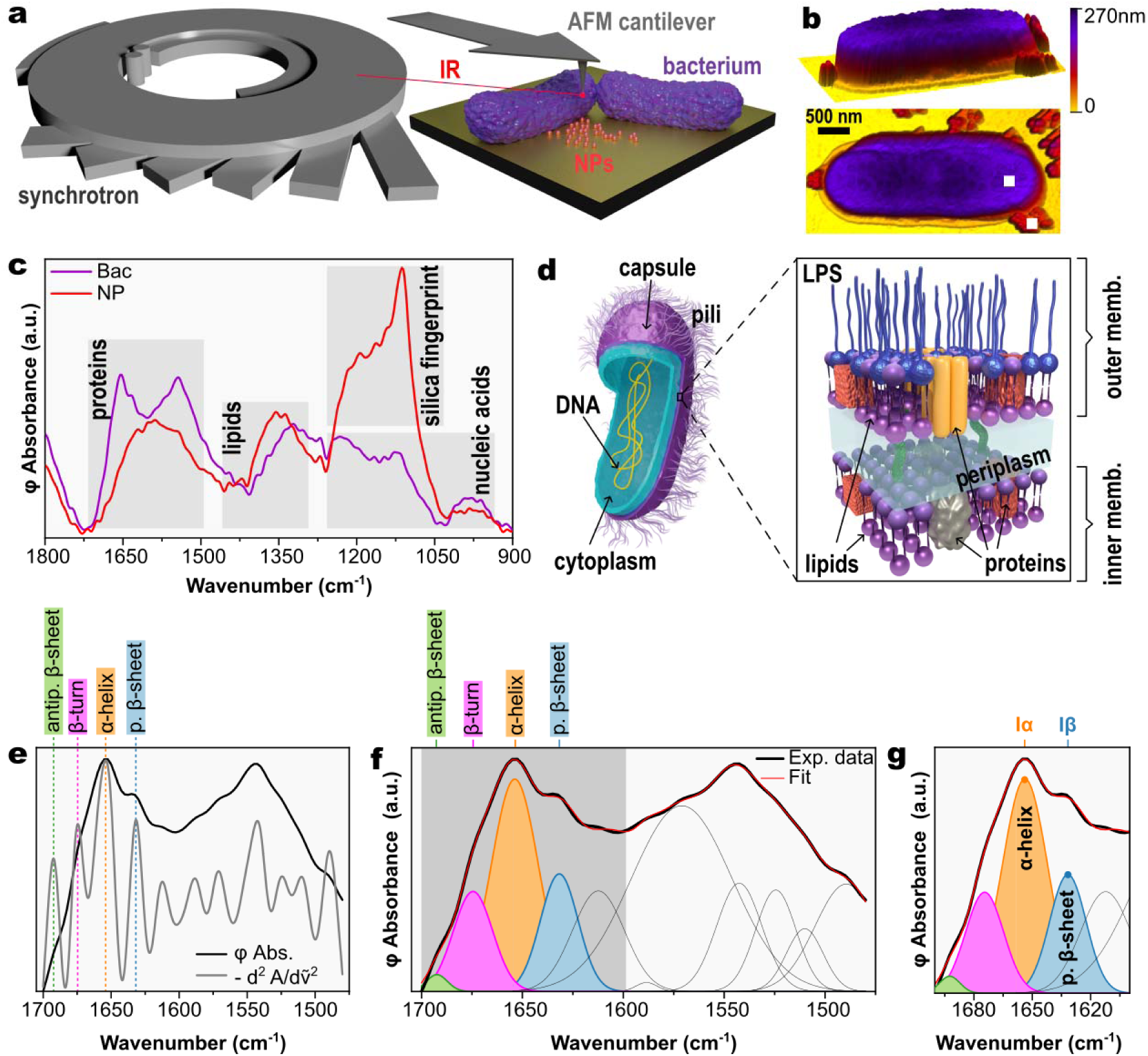
Nano-FTIR experimental workflow and spectral analysis strategy. (a) Schematic illustration of the nano-FTIR experiment performed at the Imbuia beamline, SIRIUS, Campinas, Brazil. (b) AFM image of *E. coli* incubated with bare-SiO_2_ (scale bar, 500 nm). The two marked positions indicate the locations where IR spectra were acquired. (c) IR spectra collected at the positions highlighted in (b), with characteristic biochemical fingerprint bands indicated. (d) Schematic representation of the internal structure of a Gram-negative bacterium and an enlarged view of membrane components. (e) Normalized IR absorbance spectra in the Amide I and Amide II regions together with their normalized negative second-derivative spectra. (f) Gaussian peak fitting of the spectral region shown in (e), with peak positions defined from the negative second-derivative spectra. (g) Gaussian components assigned to α-helical and β-sheet structures were used to determine the α-helix (Iα) and β-sheet (Iβ) intensities, from which the Iα/Iβ ratio was calculated.

Fig. 4b shows an AFM image of *E. coli* incubated with bare-SiO_2_, where rod-shaped bacteria are observed alongside nanoparticle aggregates, consistent with SEM observations (Fig. 2a). Two representative locations are highlighted in the topography map, one on the bacterial cell and another on the nanoparticle aggregate. Fig. 4c displays the corresponding SINS phase spectra acquired at these positions. The spectrum collected on the bacterial region (purple curve) exhibits characteristic infrared bands of *E. coli*, including contributions from proteins, lipids, nucleic acids, and carbohydrates, in agreement with previous reports ^22–24^. In particular, bands at approximately 1655 and 1545 cm^-1^ correspond to the amide I and amide II modes, respectively, which are commonly used as markers of protein secondary structure ^55–59^. Additional features between 1350 and 1000 cm^-1^ arise predominantly from phosphodiester vibrations of nucleic acids and phospholipids, as well as carbohydrate-related C–O–C and P–O–C modes ^22–24^. A schematic representation of the GN bacterial envelope, highlighting the OM, periplasmic space, and inner membrane, is provided in Fig. 4d ^60,61^.

In contrast, the spectrum acquired on the nanoparticle aggregate (red curve in Fig. 4c) shows the characteristic silica fingerprint between 1200 and 1050 cm^-1^, with no detectable bacterial contribution ^17^. To enable quantitative comparison across samples, all SINS spectra were processed using an identical workflow. First, the 1480–1700 cm^-1^ region, encompassing the amide I and amide II bands, was min–max normalized, followed by calculation of the negative second derivative using a five-point Savitzky–Golay filter (window = 5 points, 12.5 cm^-1^; polynomial order = 3). The filter was applied directly to the normalized spectrum without prior smoothing, following the approach of Yang et al. (Fig. 4e) ^59^. The maxima of the negative second derivative were used as fixed center positions for Gaussian components in the subsequent fitting of the unsmoothed experimental data (Fig. 4f). This strategy minimizes parameter correlation associated with strongly overlapping amide bands, stabilizes nonlinear convergence, and improves the reproducibility of comparative spectral analysis across samples ^59,62,63^.

Four Gaussian components associated with amide I secondary-structure contributions were identified for the *E. coli* control sample ^55–58,64^, and the ratio between the amplitudes of the α-helical (∼1655 cm^-1^) and parallel β-sheet (∼1635 cm^-1^) components (Iα/Iβ), the two principal protein secondary structures, was used as a comparative descriptor across samples (Fig. 4g). Peak amplitudes were used rather than integrated areas, as they are directly optimized during fitting and exhibit lower uncertainty propagation under strong band overlap conditions. Fitting parameters and corresponding Iα/Iβ ratios obtained for spectra within the same sample group were highly consistent, confirming the reproducibility of the analysis (Fig. S8–S9 and Section S5 in the Supporting Information).

Peak intensity uncertainties and the uncertainty of the Iα/Iβ ratio were estimated by residual bootstrap resampling (B = 300), with error propagation performed within each iteration to capture inter-component covariance ^65^. In nonlinear fitting, uncertainties derived from the linearized covariance matrix can be underestimated, particularly when parameter correlations and strong nonlinearity are present. Bootstrap resampling provides a model-independent alternative that can yield more realistic confidence intervals ^66,67^. Here, due to the overlapping and correlated nature of the amide I and II Gaussian components, bootstrap was used to obtain more reliable estimates of parameter variability. All corresponding datasets are provided in the Supporting Information.

Fig. 5a–e presents AFM images of representative samples, including control *E. coli* and *E. coli* incubated with bare-, man-, mal-, and tre-SiO_2_, respectively. In each image, the white square indicates the region selected for SINS spectral acquisition on the bacterial cell (measurements performed at the bacterium–nanoparticle interface and directly on nanoparticles are provided in Section S4 of the Supporting Information). The corresponding infrared spectra are shown in Fig. 5f, and the associated second-derivative analysis is presented in Fig. S14. Identical data-processing procedures were applied to all samples (Fig. S8–S13 at Supporting Information), and the Iα/Iβ ratio was compared across the five conditions in Fig. 5g, along with its associated uncertainty (σ_R_), estimated by residual bootstrap resampling to account for the covariance between σ(Iα) and σ(Iβ). Error bars represent ±2σ, corresponding to the 95% confidence interval.

**Figure 5.**
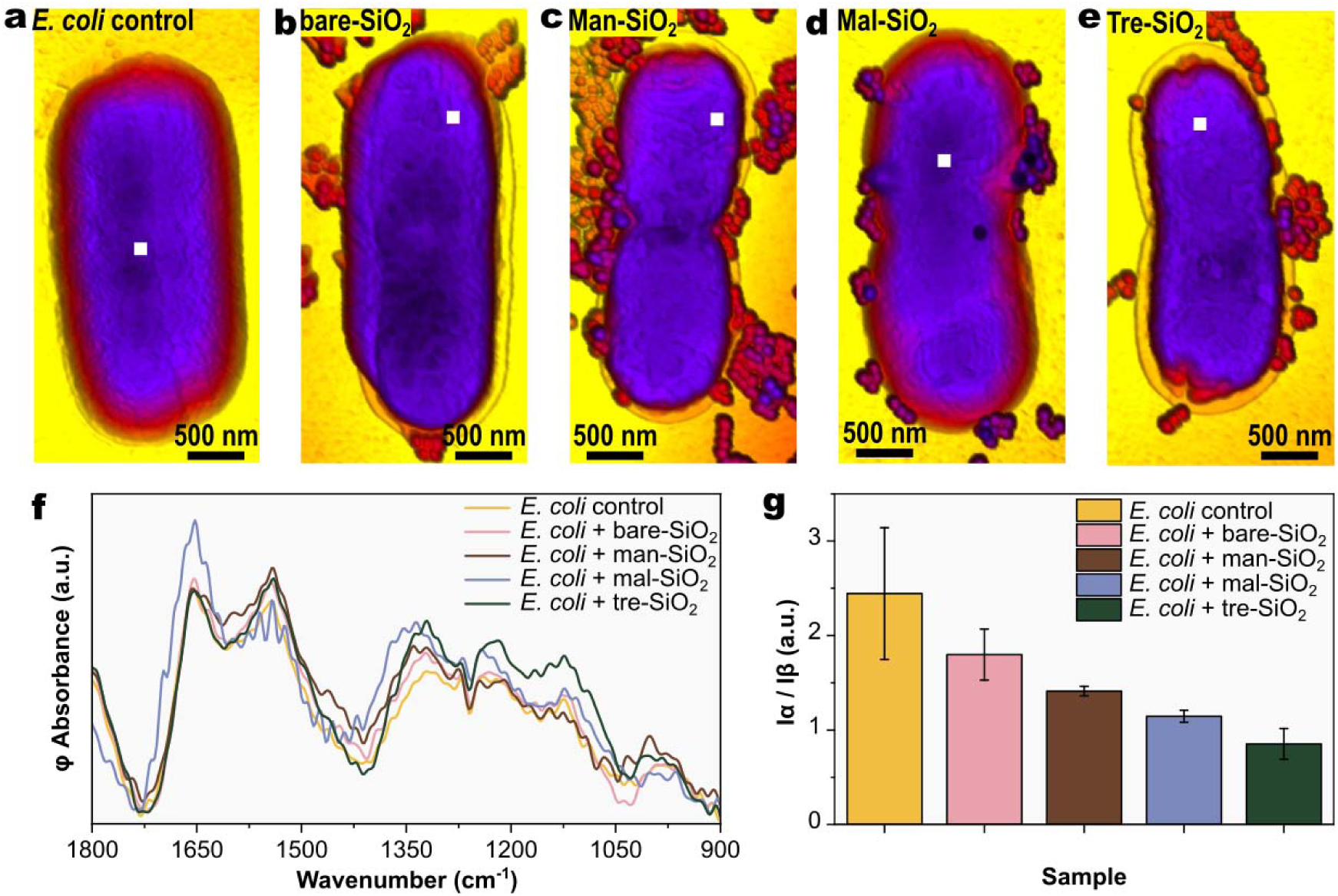
Nano-FTIR analysis of *E. coli* incubated with NPs. (a–e) AFM images of control *E. coli* and cells incubated with bare-SiO_2_, man-SiO_2_, mal-SiO_2_, and tre-SiO_2_, respectively. The marked positions indicate the locations where nano-FTIR spectra were acquired. (f) Nano-FTIR absorption spectra collected at the positions indicated in (a–e). (g) Comparison of the Iα/Iβ ratios obtained for each sample. Error bars represent the propagated uncertainty estimated by residual bootstrap resampling (B = 300 iterations), accounting for the covariance between the uncertainties associated with Iα and Iβ (±2σ, corresponding to a 95% confidence interval).

This comparison was designed to assess whether nanoparticle incubation induces detectable spectral changes in bacterial cells, even in the absence of nanoparticles directly observed at the measurement location. A systematic decrease in the Iα/Iβ ratio was observed for *E. coli* incubated with carbohydrate-functionalized nanoparticles relative to the control and bare-SiO_2_ conditions, indicating, within the applied spectral model, an increased contribution of β-sheet relative to α-helical components. This effect was more pronounced for the disaccharide-coated nanoparticles (maltose and trehalose) than for the monosaccharide-coated nanoparticles, whereas the control and bare-SiO_2_ samples exhibited comparable Iα/Iβ values. The fitted intensity values were stable across bootstrap iterations, with the associated uncertainties confirming the robustness of the Gaussian peak fitting (Tables S5–S9). Statistical comparison using a two-tailed z-test based on bootstrap-derived uncertainties further showed no significant difference between the control and bare-SiO_2_ groups at the 95% confidence level, whereas all carb-SiO_2_ differed significantly from the control and from one another (Table S10).

All SINS measurements were performed directly on bacterial cells, indicating that the observed spectral differences arise from nanoparticle incubation rather than from direct nanoparticle contributions. These results demonstrate that SINS is sensitive to carbohydrate-dependent perturbations induced by nanoparticle exposure at the single-cell level.

Fig. 6 compares SINS spectra acquired directly on bacterial cells with those collected at the bacterium–nanoparticle interface for *E. coli* incubated with bare-, man-, mal-, and tre-SiO_2_. A schematic representation of the two measurement locations is shown in Fig. 6a, and the corresponding Iα/Iβ ratios, obtained following identical data processing, are summarized in Fig. 6b. As discussed above, incubation with carbohydrate-functionalized nanoparticles shifts the protein secondary structure balance toward a greater relative contribution of β-sheet compared with α-helical conformations. Notably, the difference in the Iα/Iβ ratio between measurements acquired on the bacterial surface and at the bacterium–nanoparticle interface progressively decreased from approximately 33% for bare-SiO_2_ to 21% for man-SiO_2_ and mal-SiO_2_, reaching only 2% for tre-SiO_2_. This trend suggests that carbohydrate-functionalized nanoparticles induce structural perturbations that extend beyond the bacterium–nanoparticle interface, whereas bare-SiO_2_ produces only localized perturbations that remain largely confined to the contact region.

**Figure 6.**
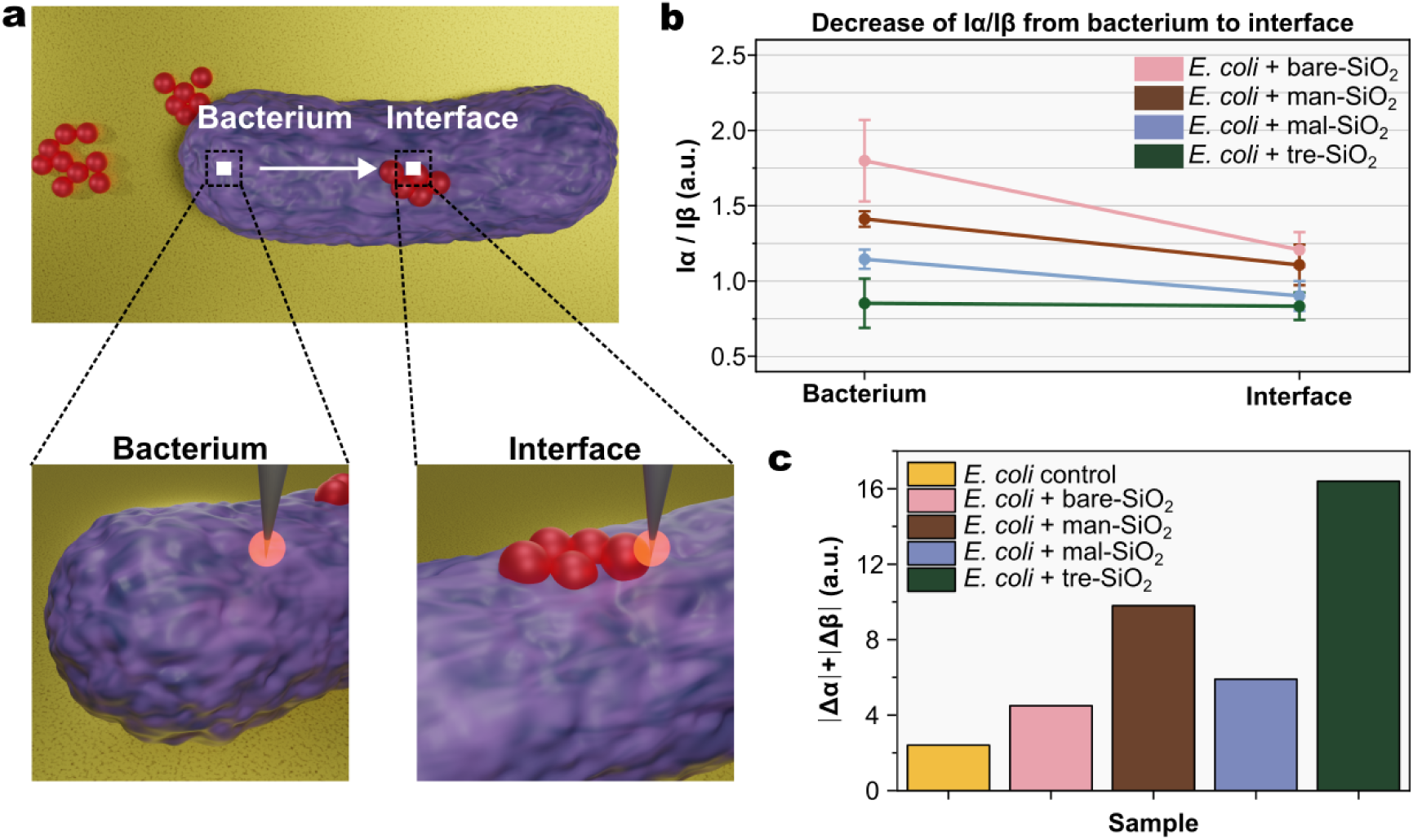
Comparison of nano-FTIR measurements on bacterial cells and at the bacterium–nanoparticle interface. (a) Schematic illustration of the measurement strategy, showing the AFM tip positioned above the bacterium and subsequently at the bacterium–nanoparticle interface. (b) Comparison of the Iα/Iβ ratios obtained at the two measurement locations for each sample, comparing measurements acquired on the bacterial surface and at the bacterium–nanoparticle interface. Error bars represent the propagated uncertainty estimated by residual bootstrap resampling (B = 300 iterations), accounting for the covariance between the uncertainties associated with Iα and Iβ (±2σ, corresponding to a 95% CI). (c) Cumulative absolute shift of the α-helical and β-sheet bands (|Δα-helix| + |Δβ-sheet|), calculated from the differences in the positions of the α-helical (≈1656 cm^-1^) and β-sheet (≈1635 cm^-1^) bands between spectra acquired on the bacterial surface and at the bacterium–nanoparticle interface for each sample. The control *E. coli* was included as a reference, with the peak shifts calculated between two different measurement locations within the same control group.

To further evaluate these confined interfacial spectral perturbations, the positions of the α-helical and β-sheet bands were extracted, and the cumulative absolute peak shift (|Δα-helix| + |Δβ-sheet|) was calculated from the differences in band positions between spectra acquired on the bacterial surface and at the bacterium–nanoparticle interface. Because the amide I band is predominantly sensitive to protein backbone secondary structure, with minimal contributions from side-chain vibrations ^56^, shifts in these bands reflect changes in the local protein conformational environment. As shown in Fig. 6c, the cumulative absolute peak shift increased for carbohydrate-functionalized nanoparticles, with tre-SiO_2_ exhibiting the largest spectral perturbation. Error bars are not shown because the cumulative absolute peak shift itself is the reported metric. As a reference, the control *E. coli* sample gave a cumulative absolute peak shift of 2.4 cm^-1^, while bare-, man-, mal-, and tre-SiO_2_ exhibited cumulative absolute peak shifts of 4.5, 9.8, 5.9, and 16.4 cm^-1^, respectively.

Notably, *E. coli* incubated with tre-SiO_2_, which exhibited the largest conformational perturbation among all samples, also showed a distinct spectral feature. SINS spectra acquired on the bacterial surface consistently revealed an additional band at approximately 1650 cm^-1^, assigned to random coil structures ^55,59,62,68^ (Fig. S13, Supporting Information). The presence of this band was confirmed in multiple independent measurements acquired on bacterial cells from the same sample, providing additional evidence that tre-SiO_2_ induces a greater disruption of the native protein secondary structure within the bacterial outer membrane.

Collectively, these results demonstrate that carbohydrate coating modulates protein conformational changes at the nano–bio interface in a manner that depends on the glycan chemistry of the nanoparticle surface. The pronounced spectral differences observed between measurements acquired on the bacterial surface and at the bacterium–nanoparticle interface indicate that nanoparticle association creates a chemically distinct interfacial environment, where protein conformational changes differ from those detected elsewhere on the bacterial envelope. Because SINS probes a nanometric interaction volume beneath the AFM tip apex, it enables direct interrogation of these confined interfacial regions without averaging their spectral signatures with those of the surrounding sample ^29^. Consequently, localized changes in the Amide I profile, including variations in the Iα/Iβ ratio, peak positions, and the emergence of random coil structures, can be resolved with spatial specificity. These findings highlight the unique capability of SINS to reveal chemically confined nano-bio interactions that remain largely inaccessible to diffraction-limited vibrational spectroscopies and establish nanoscale infrared spectroscopy as a powerful tool for investigating nanoparticle-induced molecular transformations at the single-cell level.

## CONCLUSIONS

This work demonstrates that carbohydrate surface chemistry governs protein conformational remodeling at the nano–bio interface. By combining correlative electron microscopy with SINS spectroscopy, we show that these molecular responses originate at the bacterium–nanoparticle interface and can propagate across the bacterial envelope depending on nanoparticle surface chemistry. While electron microscopy established the extent of nanoparticle association with *Escherichia coli*, SINS revealed localized biochemical perturbations that cannot be inferred from morphology alone. These perturbations were characterized by systematic changes in the relative α-helical and β-sheet contributions of the amide I band, interface-dependent shifts in characteristic vibrational bands, and, in the case of tre-SiO_2_, the emergence of a random coil component, indicating increased disruption of the native protein secondary structure. Comparison of spectra acquired directly on bacterial cells and at the bacterium–nanoparticle interface further revealed that carbohydrate coating modulates both the magnitude and the spatial extent of these molecular responses, underscoring the critical role of nanoparticle surface chemistry in shaping localized protein conformational remodeling. Beyond these biological findings, this study highlights the unique capability of SINS to directly probe chemically confined molecular responses within nano–bio interfaces that remain largely inaccessible to diffraction-limited vibrational spectroscopies. The combination of nanoscale infrared spectroscopy with a robust spectral analysis framework provides a broadly applicable strategy for investigating molecular transformations at complex biological interfaces. We anticipate that this approach will facilitate mechanistic studies of nano–bio interactions and support the rational design of functional nanomaterials for biomedical and biotechnological applications.

## METHODS

### Chemicals and biological components

Tetraethoxysilane (TEOS, reagent grade 98%), ammonium hydroxide aqueous solution (NH_4_OH, 28−30% wt NH_3_, ACS reagent), ethanol (P.A.), D (+) mannose (≥99%), D (+) maltose (≥99%), D (+) trehalose (≥99%), phosphate buffer saline (PBS) tablets, bovine serum albumin (BSA), glutaraldehyde and sodium cacodylate trihydrate were purchased from Sigma-Aldrich (St. Louis, MO, USA). Sulfuric acid (H_2_SO_4_ 98%) and sodium chloride (P.A.) were acquired from Synth (SP, Brazil). Phenol liquid (90%) was obtained from Dinâmica (SC, Brazil). Water used in all procedures was obtained from a water purification system (Purelab from ELGA-BUX, U.K.) and had a measured resistivity of 18.2 MΩ·cm.

### Synthesis of Nanoparticles (NPs)

Silica nanoparticles (SiO_2_) were synthesized using the well-known Stöber process with some modifications ^69–71^. Briefly, 5.2 mL of NH_4_OH was added to 120 mL of ethanol under magnetic stirring. Two aliquots of 2.5 mL of tetraethyl orthosilicate (TEOS) were added after 0.5 and 3 h of stirring. The reaction to form SiO_2_ occurred under room temperature and constant stirring overnight. The purification of the obtained suspension was conducted through centrifugation steps - 10.000 rpm for 15 min at 4°C – 2 times in ethanol and 3 times in water. The final suspension was stored in water at 4-8°C.

The carbohydrates (mannose, maltose, and trehalose) nanoparticles (carb-SiO_2_) were obtained by physical adsorption. The carbohydrates were incubated with previously synthesized SiO_2_ in water at 37°C and 1 h under agitation with mass ratios of 1mg mL^-1^ of SiO_2_ with 5 mg mL^-1^ of carbohydrates. Carbohydrate excess was removed by centrifugation (12.000 rpm, 15 minutes) twice. The characterization and other assays conducted with carb-SiO_2_ were performed with fresh prepared particles once they were not stored.

### Characterization of SiO_2_ and carb-SiO_2_

Stock suspension of SiO_2_ concentration was calculated by gravimetric analysis. A known suspension volume was dried at 100°C oven overnight, and the final solid was weighed. The experiment was made in triplicate.

Once the concentration was determined, the hydrodynamic (D_H_) diameter of a 1 mg ml^-1^ suspension was measured by dynamic light scattering (DLS) using a Malvern Zetasizer ZS equipment equipped with a red laser (632.8 nm) and operated in backscatter mode (detection angle = 173°). The measurements were performed in triplicate, each consisting of 10 runs of 10 s at 37°C with thermal stabilization of 120 s. The correlation curves were analyzed using the method of cumulants to obtain the hydrodynamic diameter (Z-average) and the polydispersity index (PDI) and a nonnegative least-squares adjustment algorithm (NNLS) to extract size distributions. Both procedures are implemented in the Malvern′s Zetasizer software. The D_H_ of carb-SiO_2_ was also measured by the aforementioned method.

Morphology and size distribution of bare-SiO_2_ and carb-SiO_2_ were evaluated by scanning transmission electron microscopy (STEM). The samples were prepared by depositing 10 μL of the suspensions on a 300-mesh copper grid (TedPella) with support carbon film. Samples were examined using a SEM-FEI Inspect F50 microscope with an acceleration voltage of 30 kV. The experiments were carried out at the Brazilian Nanotechnology Laboratory (LNNano) electron microscopy facility. Data processing was performed using ImageJ software. To obtain size distribution, at least 200 nanoparticles were counted.

After carbohydrates incubation, the sugars removal were evaluated by the consolidated phenol-sulfuric acid method proposed by Dubois (1956). Basically, during 4 centrifugation steps, the supernatants were collected and had the total sugar quantification. After the second centrifugation, no considerable sugar was detected on the supernatant once all carbohydrate excess had been removed.

The evaluation of parameters such as morphology, size and presence of carbohydrates on the surface of SiO_2_ were carried out using the Transmission Electron Microscopy (TEM) technique in the Brazilian Nanotechnology Laboratory (LNNano). The microscope used was the JEOL JEM2100F operated at 200 kV and equipped with a Gata Image Filter (GIF) spectrometer, which allowed obtaining filtered images in different energy ranges (EFTEM) and an Orisis camera for obtaining conventional TEM images (CTEM). The samples were prepared by deep coating a 400 Mesh copper grid (TedPella) containing holey-carbon in the freshly prepared NPs suspension (concentration of 1 mg mL^-1^), followed by drying at room temperature. For EFTEM analysis, two types of images were obtained in two different modes, zero-loss and core-loss. The EFTEM zero-loss analyzes were obtained by applying a filter in the region of the zero-loss peak and thus allowed obtaining images with greater contrasts. In contrast, the EFTEM core-loss analyzes are obtained by applying a filter to the silicon and carbon energies and obtaining images for each of these filters. The result is to obtain a compositional map of the samples and thus identify exactly the presence of carbon from carbohydrates on the surface of the SiO_2_.

### Interaction between E. coli and carb-SiO_2_

Bacterial experiments followed previously established procedures outlined in our prior publication ^17^. In summary, an *E. coli* (number 5880, André Tosello Foundation, Campinas – Brazil) suspension was derived from cryopreserved stock colonies, sustained in Luria Bertani broth (LB, containing 10 g L^−1^ of peptone and NaCl, and 5 g L^−1^ of yeast extract, from Himedia Laboratories) using an orbital shaker set at 37°C and 250 rpm. Following cultivation, the bacterial solution underwent purification through centrifugation (4000 rpm, 5 min) and subsequent resuspension in a saline solution (0.9% NaCl) repeated twice. Subsequently, the cell concentration was determined by measuring OD600.

For the bacterial susceptibility assay, 200 μL of *E. coli* at a concentration of 10^6^ colony forming units (CFU mL^−1^) and varying concentrations of NPs (62.5 to 1,000 μg mL^-1^) were co-incubated in a 96-well plate at 37°C. After 24 hours, each well absorbance at λ=600 nm was recorded in a Perkin Elmer EnSpire® 2300 plate reader, and bacterial growth was quantified, with 100% growth referenced to the negative control (*E. coli* in LB broth). A positive control containing 4 μg mL^-1^ ampicillin and blank plates with pure LB were included. The optical influence of NPs was excluded by subtracting the absorbance value of pure NPs incubated with LB, referred to as “blank NPs”. The experiments were conducted with triplicates and repeated on three different days (N=3). Results were statistically analyzed in GraphPad Prism software with two-way ANOVA.

For the analysis of the interaction between NPs and the bacterial OM, samples were prepared for scanning electron microscopy (SEM). A suspension containing 10^9^ CFU mL^−1^ of *E. coli* and 1 mg mL^−1^ of bare-SiO_2_ and carb-SiO_2_ was dispersed in LB medium. The resulting mixture was incubated in an orbital shaker (37 °C, 250 rpm) for 15 minutes. Following this incubation period, bacteria were collected and purified with PBS through centrifugation (4000 rpm, 5 min) repeated twice. The final pellet was resuspended in PBS to obtain a highly concentrated bacterial suspension, which was deposited onto a glass substrate previously treated with 0.01% poly-L-lysine (PLL). Once the bacterial adhered to the modified substrate, a 3% glutaraldehyde in sodium cacodylate buffer 0.1M was introduced into a device containing the substrate, ensuring the sample remained submerged overnight. After the fixation process, the glutaraldehyde solution was replaced with cacodylate buffer 0.1M and exchanged three times in the device. Finally, the samples were gradually dehydrated using ascending concentrations of ethanol (15%, 30%, 50%, 70%, 80%, 90%, and 100%, 15 minutes each). The dehydrated samples were then subjected to critical point drying (CPD) of the carbon dioxide (CO_2_) in a Bal Tec CPD 030 (Bal-Tec-Liechtenstein). Analysis of the samples was conducted using SEM-FEI Inspect F50 (LNNano) with a tension of 2 kV. For transmission electron microscopy (TEM), the preparation process was like SEM, but with some differences. The samples were pelleted and centrifuged after each solution change. An additional post-fixation and staining step with 1% osmium tetroxide reduced with 1.5% potassium ferrocyanide in 0.1M sodium cacodylate buffer was performed prior to dehydration. After dehydration, the samples were infiltrated in epoxy resin (Embed812) and polymerized for 48 h at 60°C. Ultrathin sections (70–100 nm) were obtained by ultramicrotomy and analyzed in a JEOL JEM-2100 transmission electron microscope at 80 keV.

For SINS analysis at the Imbuia beamline at SIRIUS (the 4^th^ generation Synchrotron source in Campinas, Brazil) ^32^, samples were prepared following the same incubation procedure described for SEM. To remove LB, saline solution was employed since PBS interferes with IR bands. The final pellet was resuspended in autoclaved water and then deposited onto a gold-Si substrate for subsequent analysis.

Atomic force microscopy (AFM) images were acquired at the Imbuia beamline to guide the SINS experiments. Initial topography images were obtained using a metallic AFM tip, which provides superior resolution. The same sample regions were subsequently imaged with the Nano-FTIR tip. Since both sets of images correspond to the same regions, only the AFM-tip images are presented here, as they exhibit higher quality.

All SINS spectra were collected with a spectral resolution of 10 cm^-1^, using 30 averages per spectrum and an integration time of 20.1 ms. Phase adjustment and baseline alignment were performed with NeaPlot software. Spectra were analyzed in the phase representation at the second harmonic.

For data analysis, three spectral regions were selected: I) 1700–1480 cm^-1^, corresponding to the amide I and amide II bands; II) 1480–1300 cm^-1^, associated with lipid vibrational modes; and III) 1260–1020 cm^-1^, which provides information on nucleic acids in bacterial samples. Regions II and III were analyzed by negative second-derivative processing for qualitative band identification only, without Gaussian decomposition or quantitative fitting. The detailed Gaussian decomposition and statistical analysis described below apply exclusively to region I.

For protein secondary structure analysis, the 1480–1700 cm^-1^ region was processed as follows. Each spectrum was normalized to the [0, 1] range using min-max scaling. The negative second derivative was computed using a five-point Savitzky–Golay filter (window = 5 points, 12.5 cm^-1^ coverage; polynomial order = 3) applied to the normalized spectrum without prior smoothing, following the protocol of Yang et al ^59^. No baseline correction was applied to the second-derivative spectra prior to peak detection; the raw second derivative was used directly. Peak positions were identified as maxima of the negative second derivative above an adaptive threshold. In cases where a spectral component was visible in the raw second derivative but fell below the automatic detection threshold — typically due to truncation at the spectral window boundary or low signal-to-noise at the window edges — the corresponding center frequency was added manually after visual inspection of the derivative profile and verified by improvement in the RMSE of the subsequent Gaussian fit.

Gaussian decomposition was performed jointly over the amide I and amide II region (1480–1700 cm^-1^) rather than over the amide I band alone. This choice was motivated by the presence of a residual water absorption band in the 1580–1620 cm^-1^ transition zone between the two amide bands, which creates a spectral continuum linking them. Restricting the decomposition to the amide I region (1600–1700 cm^-1^) resulted in poor boundary fitting and systematic overestimation of the amide I Gaussian amplitudes, as components at the low-wavenumber edge of the amide I band expanded artificially to compensate for the unmodelled water contribution. Including the amide II region in the fitting window provided the necessary spectral context for the boundary Gaussians, yielding more stable convergence and physically consistent amplitude estimates for all amide I components.

Gaussian decomposition was performed on the unsmoothed normalized spectra, with peak centers fixed at the positions identified from the second derivative, while amplitudes and widths were optimized freely within physically motivated bounds (amplitude: 0.01–1.5; width: 5–25 cm^-1^). The fitting was implemented using the SciPy curve_fit function with the Trust Region Reflective (TRF) algorithm (maximum 20,000 function evaluations). The quality of each decomposition was assessed by the root mean square error (RMSE) between the experimental spectrum and the reconstructed curve.

Spectral decompositions were excluded from further analysis when one or more of the following criteria were met: (i) bootstrap uncertainty of the α-helix/β-sheet intensity ratio exceeding 20%; (ii) 95% confidence interval including non-physical (negative) values; (iii) RMSE > 0.016; (iv) separation between the α-helix and β-sheet Gaussian centers below 15 cm^-1^; or (v) individual Gaussian components with bootstrap uncertainty exceeding 100% of their fitted amplitude.

The α-helix/β-sheet ratio was defined as the ratio of the fitted amplitudes (peak intensities) of the Gaussian components assigned to the α-helix (∼1655 cm^-1^) and parallel β-sheet (∼1635 cm^-1^) bands within the amide I region. Peak amplitudes were preferred over integrated areas because they are directly optimized during curve fitting and exhibit lower uncertainty propagation under the strong band overlap conditions characteristic of the amide I region in near-field infrared spectra — a behavior confirmed empirically in the present dataset, where area-based ratios showed systematic instability when the β-sheet component width approached the imposed lower boundary.

Uncertainties were estimated by residual bootstrap resampling (B = 300 iterations), in which the fit residuals were resampled with replacement to generate synthetic spectra, each subsequently re-fitted under identical constraints. The standard deviation of the resulting amplitude distributions was taken as the uncertainty (σ) for each band component. The α- helix/β-sheet ratio and its uncertainty were computed within each bootstrap iteration directly from the re-fitted amplitudes, thereby implicitly accounting for the covariance between the two components. All fitting and statistical procedures were implemented in Python using a custom script. Detailed results of the Gaussian decomposition, RMSE values, fitted parameters, bootstrap uncertainty estimates, and criteria for spectral exclusion for all samples are provided in the Supporting Information.

### Cell viability

NIH-3T3 fibroblast cells were plated at a density of 10^5^ cells mL^-1^ in a 96-well plate. After 24 hours of incubation in Dulbecco’s Modified Eagle Medium (DMEM, Sigma), the cells were exposed to varying concentrations of bare-SiO_2_ and carb-SiO_2_ (62.5 to 1,000 μg mL^-1^). Following the designated incubation period (1 and 24h), cells were treated with the Alamar Blue solution (90 μL DMEM + 10 μL Alamar Blue per well) and further incubated for 4 hours at 37°C and 5% CO_2_, shielded from light. Supernatants were read on a Perkin Elmer EnSpire® 2300 plate reader using excitation at λ=560 nm and emission at λ=590 nm. Cell viability was calculated considering 100% from negative control (only cells in culture medium). Blank (only culture medium + Alamar Blue) and positive controls with known cytotoxic agents were also prepared ^72^. The experiments were conducted with triplicates and repeated on two different days (N=2). Results were statistically analyzed in GraphPad Prism software with two-way ANOVA.

## ACKNOWLEDGMENTS

The authors acknowledge the Imbuia beamline at LNLS (Proposals 20221969 and 20231377), the Electron Microscopy Laboratory at LNNano for access to electron microscopy facilities (Proposals 20221627, 20230508, 20230462, and 20240959), and the Brazilian Biosciences National Laboratory (LNBio) for providing the infrastructure required for the biological experiments. The authors thank Juliana Carvalho, Talitha Stefanello, Maiara Terra, and Flávia Galdino for technical support and valuable scientific discussions.

## AUTHOR CONTRIBUTIONS

C.L.B.F. performed the nanoparticle synthesis and physicochemical characterization, conducted the biological experiments, analyzed the data, and wrote the manuscript. A.O.P. participated in the Imbuia beamtime experiments and contributed to data acquisition. R.S.R. contributed to the preparation of biological samples for SEM and TEM analyses and assisted with data analysis. L.J.C.A. contributed to nanoparticle synthesis, physicochemical characterization, and discussion of the results. O.M.M.M.C. and R.O.F. were responsible for nano-FTIR measurements and contributed to data analysis and interpretation. L.S.C. and J.B. performed and analyzed the EF-TEM experiments. M.B.C. conceived the study, supervised the project, contributed to data interpretation, and critically revised the manuscript. All authors discussed the results, reviewed the manuscript, and approved the final version.

## REFERENCES

(1) Ahmad, M.; Khan, A. U. Global Economic Impact of Antibiotic Resistance: A Review. Journal of Global Antimicrobial Resistance. Elsevier Ltd December 1, 2019, pp 313–316. 10.1016/j.jgar.2019.05.024.

(2) Aminov, R. I. A Brief History of the Antibiotic Era: Lessons Learned and Challenges for the Future. Front. Microbiol. 2010, 1 (DEC). 10.3389/fmicb.2010.00134.

(3) Levin-Reisman, I.; Ronin, I.; Gefen, O.; Braniss, I.; Shoresh, N.; Balaban, N. Q. Antibiotic Tolerance Facilitates the Evolution of Resistance. Science (1979). 2017, 355, 826–830. 10.1126/science.aaj2191.

(4) World Health Organization. Global Action Plan on Antimicrobial Resistance; Geneva, 2015.

(5) Yeh, Y. C.; Huang, T. H.; Yang, S. C.; Chen, C. C.; Fang, J. Y. Nano-Based Drug Delivery or Targeting to Eradicate Bacteria for Infection Mitigation: A Review of Recent Advances. Front. Chem. 2020, 8 (April), 1–22. 10.3389/fchem.2020.00286.

(6) Hussain, S.; Joo, J.; Kang, J.; Kim, B.; Braun, G. B.; She, Z. G.; Kim, D.; Mann, A. P.; Mölder, T.; Teesalu, T.; Carnazza, S.; Guglielmino, S.; Sailor, M. J.; Ruoslahti, E. Antibiotic-Loaded Nanoparticles Targeted to the Site of Infection Enhance Antibacterial Efficacy. *Nat*. Biomed. Eng. 2018, 2 (2), 95–103. 10.1038/s41551-017-0187-5.

(7) Varaprasad, K.; Karthikeyan, C.; KanikiReddy, V.; Núñez, D.; Sadiku, E. R.; Briones, R. Antibiotic Nanomaterials. In Antibiotic Materials in Healthcare; Elsevier, 2020; pp 1–10. 10.1016/B978-0-12-820054-4.00001-X.

(8) Wu, S.; Huang, Y.; Yan, J.; Li, Y.; Wang, J.; Yang, Y. Y.; Yuan, P.; Ding, X. Bacterial Outer Membrane-Coated Mesoporous Silica Nanoparticles for Targeted Delivery of Antibiotic Rifampicin against Gram-Negative Bacterial Infection In Vivo. Adv. Funct. Mater. 2021, 2103442, 1–10. 10.1002/adfm.202103442.

(9) Hameed, S.; Baimanov, D.; Li, X.; Liu, K.; Wang, L. Synchrotron Radiation-Based Analysis of Interactions at the Nano-Bio Interface. Environmental Science: Nano. Royal Society of Chemistry July 13, 2022, pp 3152–3167. 10.1039/d2en00408a.

(10) Tian, X.; Chong, Y.; Ge, C. Understanding the Nano–Bio Interactions and the Corresponding Biological Responses. Frontiers in Chemistry. Frontiers Media S.A. June 10, 2020. 10.3389/fchem.2020.00446.

(11) Nikaido, H. Molecular Basis of Bacterial Outer Membrane Permeability Revisited Molecular Basis of Bacterial Outer Membrane Permeability Revisited. Microbiology and Molecular Biology Reviews 2013, 67 (4), 593–656. 10.1128/MMBR.67.4.593.

(12) Caroff, M.; Karibian, D. Structure of Bacterial Lipopolysaccharides. Carbohydr. Res. 2003, 338 (23), 2431–2447. 10.1016/j.carres.2003.07.010.

(13) Lu, H. D.; Yang, S. S.; Wilson, B. K.; McManus, S. A.; Chen, C. V. H. H.; Prud’homme, R. K. Nanoparticle Targeting of Gram-Positive and Gram-Negative Bacteria for Magnetic-Based Separations of Bacterial Pathogens. Applied Nanoscience (Switzerland) 2017, 7 (3–4), 83–93. 10.1007/s13204-017-0548-0.

(14) Xu, X. H. N.; Brownlow, W. J.; Kyriacou, S. V.; Wan, Q.; Viola, J. J. Real-Time Probing of Membrane Transport in Living Microbial Cells Using Single Nanoparticle Optics and Living Cell Imaging. Biochemistry 2004, 43 (32), 10400–10413. 10.1021/bi036231a.

(15) Kell, A. J.; Stewart, G.; Ryan, S.; Peytavi, R.; Boissinot, M.; Huletsky, A.; Bergeron, M. G.; Simard, B. Vancomycin-Modified Nanoparticles For. *ACS Nano* 2008, 2 (9), 1777–1788.

(16) Day, C. J.; Tran, E. N.; Semchenko, E. A.; Tram, G.; Hartley-Tassell, L. E.; Ng, P. S. K.; King, R. M.; Ulanovsky, R.; McAtamney, S.; Apicella, M. A.; Tiralongo, J.; Morona, R.; Korolik, V.; Jennings, M. P. Glycan:Glycan Interactions: High Affinity Biomolecular Interactions That Can Mediate Binding of Pathogenic Bacteria to Host Cells. Proc. Natl. Acad. Sci. U. S. A. 2015, 112 (52), E7266–E7275. 10.1073/pnas.1421082112.

(17) Capeletti, L. B.; de Oliveira, J. F. A.; Loiola, L. M. D.; Galdino, F. E.; da Silva Santos, D. E.; Soares, T. A.; de Oliveira Freitas, R.; Cardoso, M. B. Gram-Negative Bacteria Targeting Mediated by Carbohydrate–Carbohydrate Interactions Induced by Surface-Modified Nanoparticles. Adv. Funct. Mater. 2019, 29 (48), 1–11. 10.1002/adfm.201904216.

(18) Limqueco, E.; Passos Da Silva, D.; Reichhardt, C.; Su, F. Y.; Das, D.; Chen, J.; Srinivasan, S.; Convertine, A.; Skerrett, S. J.; Parsek, M. R.; Stayton, P. S.; Ratner, D. M. Mannose Conjugated Polymer Targeting P. Aeruginosa Biofilms. ACS Infect. Dis. 2020, 6 (11), 2866–2871. 10.1021/acsinfecdis.0c00407.

(19) Alvarez-Ordóñez, A.; Mouwen, D. J. M.; López, M.; Prieto, M. Fourier Transform Infrared Spectroscopy as a Tool to Characterize Molecular Composition and Stress Response in Foodborne Pathogenic Bacteria. Journal of Microbiological Methods. March 2011, pp 369–378. 10.1016/j.mimet.2011.01.009.

(20) Moen, B.; Janbu, A. O.; Langsrud, S.; Langsrud, Ø.; Hobman, J. L.; Constantinidou, C.; Kohler, A.; Rudi, K. Global Responses of Escherichia Coli to Adverse Conditions Determined by Microarrays and FT-IR Spectroscopy. Can. J. Microbiol. 2009, 55 (6), 714–728. 10.1139/W09-016.

(21) Faghihzadeh, F.; Anaya, N. M.; Schifman, L. A.; Oyanedel-Craver, V. Fourier Transform Infrared Spectroscopy to Assess Molecular-Level Changes in Microorganisms Exposed to Nanoparticles. Nanotechnology for Environmental Engineering 2016, 1 (1). 10.1007/s41204-016-0001-8.

(22) Saulou, C.; Jamme, F.; Girbal, L.; Maranges, C.; Fourquaux, I.; Cocaign-Bousquet, M.; Dumas, P.; Mercier-Bonin, M. Synchrotron FTIR Microspectroscopy of Escherichia Coli at Single-Cell Scale under Silver-Induced Stress Conditions. Anal. Bioanal. Chem. 2013, 405 (8), 2685–2697. 10.1007/s00216-013-6725-4.

(23) Hu, X. J.; Liu, Z. X.; Di Wang, Y.; Li, X. N.; Hu, J.; Lü, J. H. Synchrotron FTIR Spectroscopy Reveals Molecular Changes in Escherichia Coli upon Cu2+ Exposure. Nuclear Science and Techniques 2016, 27 (3). 10.1007/s41365-016-0067-9.

(24) Sukprasert, J.; Thumanu, K.; Phung-On, I.; Jirarungsatean, C.; Erickson, L. E.; Tuitemwong, P.; Tuitemwong, K. Synchrotron FTIR Light Reveals Signal Changes of Biofunctionalized Magnetic Nanoparticle Attachment on Salmonella Sp. J. Nanomater. 2020, 2020. 10.1155/2020/6149713.

(25) Holman, H. Y. N.; Bechtel, H. A.; Hao, Z.; Martin, M. C. Synchrotron IR Spectromicroscopy: Chemistry of Living Cells. Anal. Chem. 2010, 82 (21), 8757–8765. 10.1021/ac100991d.

(26) Holman, H. Y. N.; Miles, R.; Hao, Z.; Wozei, E.; Anderson, L. M.; Yang, H. Real-Time Chemical Imaging of Bacterial Activity in Biofilms Using Open-Channel Microfluidics and Synchrotron FTIR Spectromicroscopy. Anal. Chem. 2009, 81 (20), 8564–8570. 10.1021/ac9015424.

(27) Stiegler, J. M.; Abate, Y.; Cvitkovic, A.; Romanyuk, Y. E.; Huber, A. J.; Leone, S. R.; Hillenbrand, R. Nanoscale Infrared Absorption Spectroscopy of Individual Nanoparticles Enabled by Scattering-Type near-Field Microscopy. ACS Nano 2011, 5 (8), 6494–6499. 10.1021/nn2017638.

(28) Huth, F.; Govyadinov, A.; Amarie, S.; Nuansing, W.; Keilmann, F.; Hillenbrand, R. Nano-FTIR Absorption Spectroscopy of Molecular Fingerprints at 20 Nm Spatial Resolution. Nano Lett. 2012, 12 (8), 3973–3978. 10.1021/nl301159v.

(29) Hillenbrand, R.; Abate, Y.; Liu, M.; Chen, X.; Basov, D. N. Visible-to-THz near-Field Nanoscopy. Nat. Rev. Mater. 2025, 10 (4), 285–310. 10.1038/s41578-024-00761-3.

(30) Keilmann, F.; Hillenbrand, R. Near-Field Microscopy by Elastic Light Scattering from a Tip. *Philosophical Transactions of the Royal Society A: Mathematical*, Physical and Engineering Sciences 2004, 362 (1817), 787–805. 10.1098/rsta.2003.1347.

(31) Hermann, P.; Hoehl, A.; Patoka, P.; Huth, F.; Rühl, E.; Ulm, G. Near-Field Imaging and Nano-Fourier-Transform Infrared Spectroscopy Using Broadband Synchrotron Radiation. Opt. Express 2013, 21 (3), 2913–2919. 10.1364/OE.21.002913.

(32) Freitas, R. O.; Deneke, C.; Maia, F. C. B.; Medeiros, H. G.; Moreno, T.; Dumas, P.; Petroff, Y.; Westfahl, H. Low-Aberration Beamline Optics for Synchrotron Infrared Nanospectroscopy. Opt. Express 2018, 26 (9), 11238–11249. 10.1364/oe.26.011238.

(33) Bechtel, H. A.; Muller, E. A.; Olmon, R. L.; Martin, M. C.; Raschke, M. B. Ultrabroadband Infrared Nanospectroscopic Imaging. Proc. Natl. Acad. Sci. U. S. A. 2014, 111 (20), 7191–7196. 10.1073/pnas.1400502111.

(34) Stöber, W.; Fink, A.; Bohn, E. Controlled Growth of Monodisperse Silica Spheres in the Micron Size Range. J. Colloid Interface Sci. 1968, 69, 62–69. 10.1016/0021-9797(68)90272-5.

(35) Dubois, M.; Gilles, K. A.; Hamilton, J. K.; Rebers, P. A.; Smith, F. Colorimetric Method for Determination of Sugars and Related Substances. Anal. Chem. 1956, 28 (3), 350–356. 10.1021/ac60111a017.

(36) Verbeeck, J.; Van Dyck, D.; Van Tendeloo, G. Energy-Filtered Transmission Electron Microscopy: An Overview. In Spectrochimica Acta - Part B Atomic Spectroscopy; 2004; Vol. 59, pp 1529–1534. 10.1016/j.sab.2004.03.020.

(37) Hofer, F.; Grogger, W.; Kothleitner, G.; Warbichler, P. *Ultramicroscopy Quantitative Analysis of EFTEM Elemental Distribution Images*; 1997; Vol. 67.

(38) Liyanage, S. H.; Yan, M. Maltose-Derivatized Fluorescence Turn-On Imaging Probe for Bacteria Detection. ACS Infect. Dis. 2023, 9 (12), 2560–2571. 10.1021/acsinfecdis.3c00403.

(39) Dippel, R.; Boos, W. The Maltodextrin System of Escherichia Coli: Metabolism and Transport. J. Bacteriol. 2005, 187 (24), 8322–8331. 10.1128/JB.187.24.8322-8331.2005.

(40) Liu, M.; Li, J.; Li, B. Mannose-Modificated Polyethylenimine: A Specific and Effective Antibacterial Agent against Escherichia Coli. Langmuir 2018, 34 (4), 1574–1580. 10.1021/acs.langmuir.7b03556.

(41) Aprikian, P.; Tchesnokova, V.; Kidd, B.; Yakovenko, O.; Yarov-Yarovoy, V.; Trinchina, E.; Vogel, V.; Thomas, W.; Sokurenko, E. Interdomain Interaction in the FimH Adhesin of Escherichia Coli Regulates the Affinity to Mannose. Journal of Biological Chemistry 2007, 282 (32), 23437–23446. 10.1074/jbc.M702037200.

(42) Zhou, J.; Jayawardana, K. W.; Kong, N.; Ren, Y.; Hao, N.; Yan, M.; Ramström, O. Trehalose-Conjugated, Photofunctionalized Mesoporous Silica Nanoparticles for Efficient Delivery of Isoniazid into Mycobacteria. ACS Biomater. Sci. Eng. 2015, 1 (12), 1250–1255. 10.1021/acsbiomaterials.5b00274.

(43) Backus, K. M.; Boshoff, H. I.; Barry, C. S.; Boutureira, O.; Patel, M. K.; D’Hooge, F.; Lee, S. S.; Via, L. E.; Tahlan, K.; Barry, C. E.; Davis, B. G. Uptake of Unnatural Trehalose Analogs as a Reporter for Mycobacterium Tuberculosis. Nat. Chem. Biol. 2011, 7 (4), 228–235. 10.1038/nchembio.539.

(44) Picco, A. S.; Mondo, G. B.; Ferreira, L. F.; De Souza, E. E.; Peroni, L. A.; Cardoso, M. B. Protein Corona Meets Freeze-Drying: Overcoming the Challenges of Colloidal Stability, Toxicity, and Opsonin Adsorption. Nanoscale 2021, 13 (2), 753–762. 10.1039/d0nr06040b.

(45) Hu, Y.; Liu, X.; Liu, F.; Xie, J.; Zhu, Q.; Tan, S. Trehalose in Biomedical Cryopreservation-Properties, Mechanisms, Delivery Methods, Applications, Benefits, and Problems. ACS Biomater. Sci. Eng. 2023, 9 (3), 1190–1204. 10.1021/acsbiomaterials.2c01225.

(46) Dumat, B.; Montel, L.; Pinon, L.; Matton, P.; Cattiaux, L.; Fattaccioli, J.; Mallet, J. M. Mannose-Coated Fluorescent Lipid Microparticles for Specific Cellular Targeting and Internalization via Glycoreceptor-Induced Phagocytosis. ACS Appl. Bio Mater. 2019, 2 (11), 5118–5126. 10.1021/acsabm.9b00793.

(47) Ahire, J. H.; Chambrier, I.; Mueller, A.; Bao, Y.; Chao, Y. Synthesis of D-Mannose Capped Silicon Nanoparticles and Their Interactions with MCF-7 Human Breast Cancerous Cells. ACS Appl. Mater. Interfaces 2013, 5 (15), 7384–7391. 10.1021/am4017126.

(48) Matias, V. R. F.; Al-Amoudi, A.; Dubochet, J.; Beveridge, T. J. Cryo-Transmission Electron Microscopy of Frozen-Hydrated Sections of Escherichia Coli and Pseudomonas Aeruginosa. J. Bacteriol. 2003, 185 (20), 6112–6118. 10.1128/JB.185.20.6112-6118.2003.

(49) Beveridge, T. J. Ultrastructure, Chemistry, and Function of the Bacterial Wall. International Review of Citology 1981, 72, 229–317. 10.1016/s0074-7696(08)61198-5.

(50) Beveridge, T. J.; Graham, L. L. Surface Layers of Bacteria. Microbiol. Rev. 1991, 55 (4), 684–705. 10.1128/mr.55.4.684-705.1991.

(51) Graham, L. L.; Beveridge, T. J.; Nanninga, N. Periplasmic Space and the Concept of the Periplasm. Trends Biochem Sci. 1991. 10.1016/0968-0004(91)90135-i.

(52) Thavarajah, R.; Mudimbaimannar, V. K.; Elizabeth, J.; Rao, U. K.; Ranganathan, K. Chemical and Physical Basics of Routine Formaldehyde Fixation. Journal of Oral and Maxillofacial Pathology. September 2012, pp 400–405. 10.4103/0973-029X.102496.

(53) Szostak, R.; Silva, J. C.; Turren-Cruz, S. H.; Soares, M. M.; Freitas, R. O.; Hagfeldt, A.; Tolentino, H. C. N.; Nogueira, A. F. Nanoscale Mapping of Chemical Composition in Organic-Inorganic Hybrid Perovskite Films. Sci. Adv. 2019, 5 (10), 2–9. 10.1126/sciadv.aaw6619.

(54) Pfitzner, E.; Heberle, J. Infrared Scattering-Type Scanning near-Field Optical Microscopy of Biomembranes in Water. Journal of Physical Chemistry Letters 2020, 11 (19), 8183–8188. 10.1021/acs.jpclett.0c01769.

(55) Yan, Z.; Li, Q.; Zhang, P. Soy Protein Isolate and Glycerol Hydrogen Bonding Using Two-Dimensional Correlation (2D-COS) Attenuated Total Reflection Fourier Transform Infrared (ATR FT-IR) Spectroscopy. Appl. Spectrosc. 2017, 71 (11), 2437–2445. 10.1177/0003702817710249.

(56) Barth, A. Infrared Spectroscopy of Proteins. Biochimica et Biophysica Acta (BBA) - Bioenergetics 2007, 1767 (9), 1073–1101. 10.1016/j.bbabio.2007.06.004.

(57) Barth, A.; Zscherp, C. What Vibrations Tell Us about Proteins. Q. Rev. Biophys. 2002, 35 (4), 369–430. 10.1017/s0033583502003815.

(58) Goormaghtigh, E.; Ruysschaert, J. M.; Raussens, V. Evaluation of the Information Content in Infrared Spectra for Protein Secondary Structure Determination. Biophys. J. 2006, 90 (8), 2946–2957. 10.1529/biophysj.105.072017.

(59) Yang, H.; Yang, S.; Kong, J.; Dong, A.; Yu, S. Obtaining Information about Protein Secondary Structures in Aqueous Solution Using Fourier Transform IR Spectroscopy. Nat. Protoc. 2015, 10 (3), 382–396. 10.1038/nprot.2015.024.

(60) Choi, U.; Lee, C. R. Antimicrobial Agents That Inhibit the Outer Membrane Assembly Machines of Gram-Negative Bacteria. J. Microbiol. Biotechnol. 2019, 29 (1), 1–10. 10.4014/jmb.1804.03051.

(61) Yamanaka, M.; Hara, K.; Kudo, J. Bactericidal Actions of a Silver Ion Solution on Escherichia Coli, Studied by Energy-Filtering Transmission Electron Microscopy and Proteomic Analysis. Appl. Environ. Microbiol. 2005, 71 (11), 7589–7593. 10.1128/AEM.71.11.7589-7593.2005.

(62) Dong, A.; Huang, P.; Caughey1, W. S. Protein Secondary Structures in Water from Second-Derivative Amide I Infrared Spectra. Biochemistry 1990, 29 (13), 3303–3308. 10.1021/bi00465a022.

(63) Susi, H.; Byler, M. D. Protein Structure by Fourier Transform Infrared Spectroscopy: Second Derivative Spectra. Biochem. Biophys. Res. Commun. 1983, 115 (1), 391–397. 10.1016/0006-291X(83)91016-1.

(64) Jackson, M.; Mantsch, H. H. The Use and Misuse of FTIR Spectroscopy in the Determination of Protein Structure. Crit. Rev. Biochem. Mol. Biol. 1995, 30 (2), 95–120. 10.3109/10409239509085140.

(65) Efron, B.; Tibshirani, R. J. An Introduction to the Bootstrap; Chapman and Hall/CRC, 1994. 10.1201/9780429246593.

(66) Kazmierczak, N. P.; Chew, J. A.; Vander Griend, D. A. Bootstrap Methods for Quantifying the Uncertainty of Binding Constants in the Hard Modeling of Spectrophotometric Titration Data. Anal. Chim. Acta 2022, 1227, 1–10. 10.1016/j.aca.2022.339834.

(67) Sohn, R. A.; Menke, W. Application of Maximum Likelihood and Bootstrap Methods to Nonlinear Curve fit Problems in Geochemistry. *Geochemistry, Geophysics*, Geosystems 2002, 3 (7), 1–17. 10.1029/2001gc000253.

(68) Byler, D. M.; Susi, H. Examination of the Secondary Structure of Proteins by Deconvolved FTIR Spectra. Biopolymers 1986, 25, 469–487. 10.1002/bip.360250307.

(69) Galdino, F. E.; Picco, A. S.; Capeletti, L. B.; Bettini, J.; Cardoso, M. B. Inside the Protein Corona: From Binding Parameters to Unstained Hard and Soft Coronas Visualization. Nano Lett. 2021, 21 (19), 8250–8257. 10.1021/acs.nanolett.1c02416.

(70) Ferreira, L. F.; Picco, A. S.; Galdino, F. E.; Albuquerque, L. J. C.; Berret, J. F.; Cardoso, M. B. Nanoparticle-Protein Interaction: Demystifying the Correlation between Protein Corona and Aggregation Phenomena. ACS Appl. Mater. Interfaces 2022, 14 (25), 28559–28569. 10.1021/acsami.2c05362.

(71) Silva, C. E. P.; Picco, A. S.; Galdino, F. E.; de Burgos Martins de Azevedo, M.; Cathcarth, M.; Passos, A. R.; Cardoso, M. B. Distinguishing Protein Corona from Nanoparticle Aggregate Formation in Complex Biological Media Using X-Ray Photon Correlation Spectroscopy. Nano Lett. 2024, 24 (42), 13293–13299. 10.1021/acs.nanolett.4c03662.

(72) Sousa Ribeiro, I. R.; da Silva, R. F.; Rabelo, R. S.; Marin, T. M.; Bettini, J.; Cardoso, M. B. Flowing through Gastrointestinal Barriers with Model Nanoparticles: From Complex Fluids to Model Human Intestinal Epithelium Permeation. ACS Appl. Mater. Interfaces 2023, 15 (30), 36025–36035. 10.1021/acsami.3c07048.

