## SUPPORTING INFORMATION for "Synchrotron Nano-FTIR Reveals Carbohydrate-Dependent Protein Conformational Changes at Bacterium–Nanoparticle Interfaces"

Supporting Table of Contents

Section S1. Free carbohydrate removal ... 3

Section S2. Nanoparticles physicochemical characterization ... 4

Section S3. *In vitro* bio assays ... 10

Section S4. Nano-FTIR results ... 14

Section S5. Amide I and II band deconvolution and bootstrap uncertainty estimation 27

Section S5.1 Gaussian decomposition model ... 28

Section S5.2 Bootstrap uncertainty estimation ... 29

Section S5.3 Propagation of uncertainty for the Iα/Iβ ratio ... 30

Section S5.4 Results ... 32

Section S5.5 Statistical Comparison of Iα/Iβ Ratios Between Conditions ... 35

REFERENCES ... 38

#### Section S1. Free carbohydrate removal

The carbohydrate content in the supernatant was measured during four centrifugation steps. Following each centrifugation, the supernatant was collected and analyzed using the phenol-sulfuric method [1]. Fig. S1 displays the calculated amount of sugar in µg $ml^{-1}$. Analytical calibration curves were prepared for each sugar to correlate the absorbance values of the supernatant with sugar concentration. After the second centrifugation step, negligible sugar levels were detected in the supernatant. Therefore, a standardized procedure of two centrifugation steps was adopted to remove excess carbohydrates on the following tests.

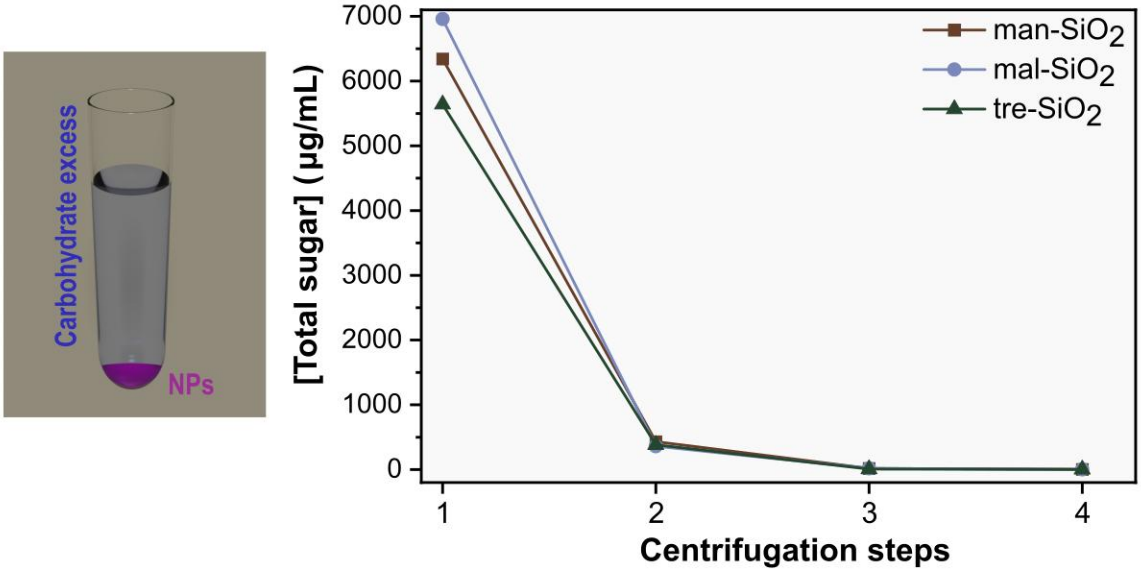


**Figure S1. Quantification of residual carbohydrates during nanoparticle purification.** Carbohydrate concentration in the supernatant determined by the phenol–sulfuric acid assay after each of the four centrifugation steps.

#### Section S2. Nanoparticles physicochemical characterization

The nanoparticles (NPs) underwent morphological characterization using Scanning Transmission Electron Microscopy (STEM). Fig. S2 illustrates the STEM images of silica nanoparticles (bare-$SiO_2$) and carbohydrates (mannose, maltose, and trehalose) nanoparticles (carb-$SiO_2$ - man-$SiO_2$, mal-$SiO_2$ and tre-$SiO_2$, respectively), revealing spherical shapes with uniform distribution across all particles. Additionally, the diameter of the NPs was measured using ImageJ software, with at least 300 particles counted for each sample. The size distribution of NPs is depicted in Fig. S2, with pink representing bare-$SiO_2$, brown representing man-$SiO_2$, blue representing mal-$SiO_2$, and green representing tre-$SiO_2$. The mean diameter of all four types of NPs was approximately 70 nm. However, it is worth noting that the carbohydrate coating was not visible using this technique.

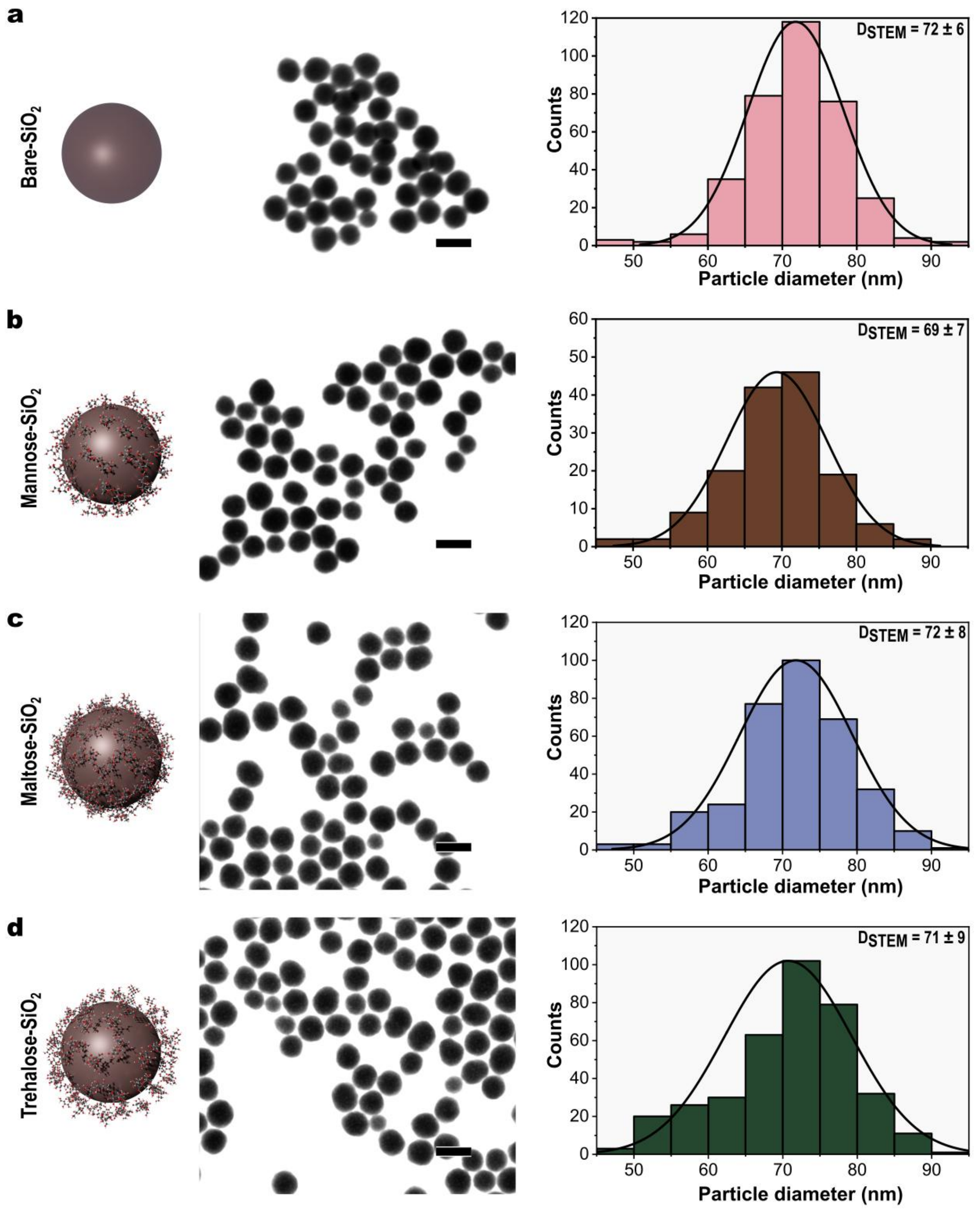


**Figure S2. STEM images and particle size distributions of NPs.** STEM images and corresponding particle size histograms obtained from ImageJ analysis for (a) bare-$SiO_2$, (b) man-$SiO_2$, (c) mal-$SiO_2$, and (d) tre-$SiO_2$.

Dynamic light scattering (DLS) was employed to measure the hydrodynamic diameter ($D_H$) of particles. Table S1 presents the mean $D_H$ values and compares them with the diameters measured using STEM ($D_{STEM}$). It is evident that the presence of carbohydrates on the silica surface increases the difference between the wet and dry diameters. This observation likely arises because DLS detects physically adsorbed carbohydrates on the nanoparticle surface. When NPs are in a biological medium, they rapidly form a biomolecular corona composed of proteins, lipids, and sugars [2,3]. While much attention has been focused on investigating the protein corona characteristics [4–7], other biomolecules such as carbohydrates can also alter the identity of NPs in a biological environment. Carbohydrate-coated NPs have been reported to enhance cellular uptake, reduce NPs toxicity, and promote cryopreservation [8–11]. DLS analysis revealed that the thickness of the carbohydrate layer increases with the length of the biomolecules. Disaccharides such as maltose and trehalose exhibited a more pronounced increase in $D_H$, with increments of 5 nm and 8 nm, respectively, compared to bare-$SiO_2$.

**Table S1. Comparison of hydrodynamic and STEM-derived diameters of NPs.** Hydrodynamic diameter, mean STEM diameter, and the difference between the two measurements.

| | **Hydrodynamic Diameter ($D_H$)** | **STEM Diameter ($D_{STEM}$)** | **$D_H$-$D_{STEM}$** | **$\Delta$($D_H$-$D_{STEM}$) (carb-$SiO_2$)-(bare-$SiO_2$)** |
|---|---|---|---|---|
| *Bare-$SiO_2$* | 108.1 ± 0.9 nm | 71.8 | 36.3 | - |
| *Man-$SiO_2$* | 107.5± 0.8 nm | 69.3 | 38.2 | 1.9 |
| *Mal-$SiO_2$* | 113.2 ± 1.1 nm | 71.8 | 41.4 | 5.1 |
| *Tre-$SiO_2$* | 116.2 ± 0.6 nm | 70.7 | 45.5 | 9.2 |

Colloidal stability is a critical parameter for biomedical nanoparticles, as aggregation can reduce cellular uptake, targeting efficiency, and drug delivery efficacy. Furthermore, particle aggregation can lead to the premature clearance of NPs from the bloodstream [12]. Figs. S3 and S4 depict the colloidal stability of NPs in water and PBS, respectively, over a 48-hour period. In both cases, all NPs formulations remained stable, as evidenced by the maintenance of the $D_H$ throughout the assay, indicating no aggregation, even under higher ionic strength conditions (PBS).

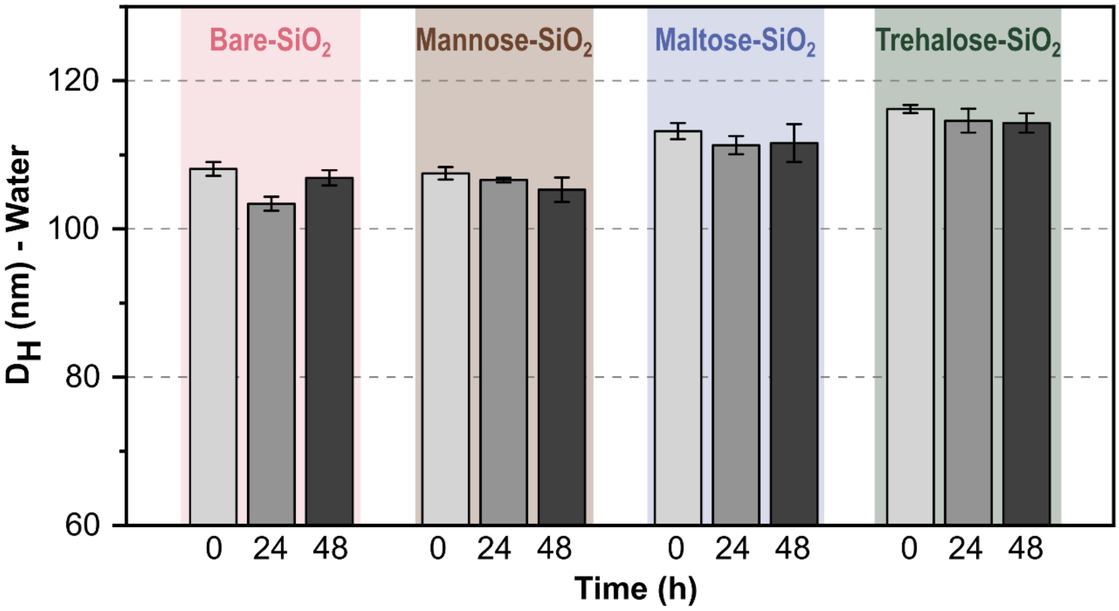


**Figure S3. Colloidal stability of NPs in water.** Hydrodynamic diameters measured by DLS at 0, 24, and 48 h after sample preparation. Pink, bare-$SiO_2$; brown, man-$SiO_2$; blue, mal-$SiO_2$; and green, tre-$SiO_2$.

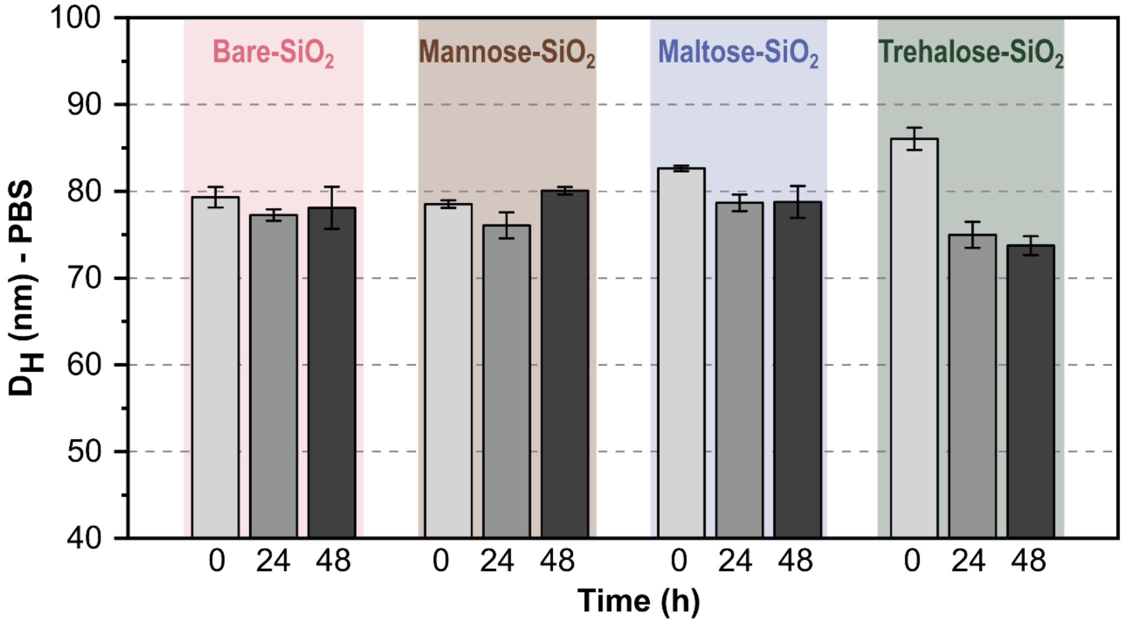


**Figure S4. Colloidal stability of NPs in PBS.** DLS measurements acquired at 0, 24, and 48 h after sample preparation. Pink, bare-$SiO_2$; brown, man-$SiO_2$; blue, mal-$SiO_2$; and green, tre-$SiO_2$.

The colloidal stability of NPs can be compromised by increasing medium complexity [13]. Fig. S5 shows the hydrodynamic diameter of NPs measured in Dulbecco's Modified Eagle Medium (DMEM), used during cell culture, and in Luria-Bertani (LB) broth, used to cultivate *E. coli*. While NPs maintained their average diameter in DMEM, LB broth induced significant particle aggregation, likely due to its high content of salts, peptides, and yeast extract.

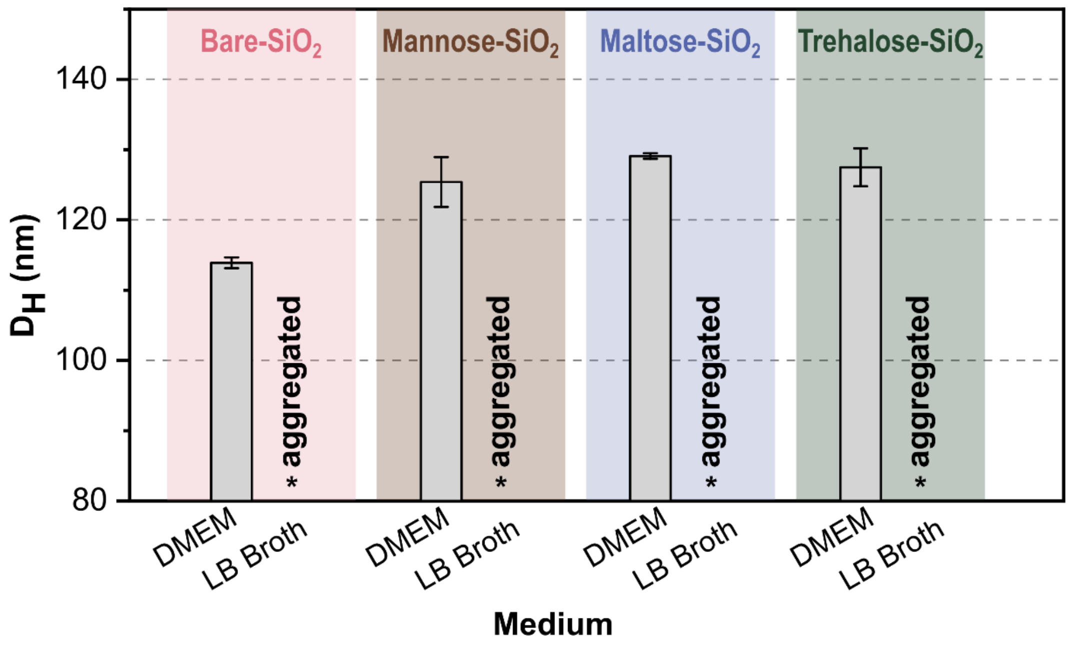


**Figure S5. Colloidal stability of NPs in DMEM and LB broth.** Hydrodynamic diameters measured by DLS in DMEM and LB broth. Pink, bare-$SiO_2$; brown, man-$SiO_2$; blue, mal-$SiO_2$; and green, tre-$SiO_2$.

#### Section S3. *In vitro* bio assays

Results of the bacterial growth assay are presented in Fig. S6, and statistical analysis of variance (Two-way ANOVA test) is described in Table S2. The growth of *E. coli* was determined by measuring absorbance at 600 nm after 24 hours of incubation with NPs. At all three concentrations tested (62.5, 125, and 250 μg $ml^{-1}$), tre-$SiO_2$ exhibited the lowest bacterial growth. Man- and mal-$SiO_2$ also impacted bacterial growth, particularly at lower concentrations. Statistical analyses were performed using a 95% confidence interval. Significance levels were defined as follows: ns, $P \geq 0.05$; *$P < 0.05$; **$P < 0.01$; ***$P < 0.001$; and ****$P < 0.0001$.

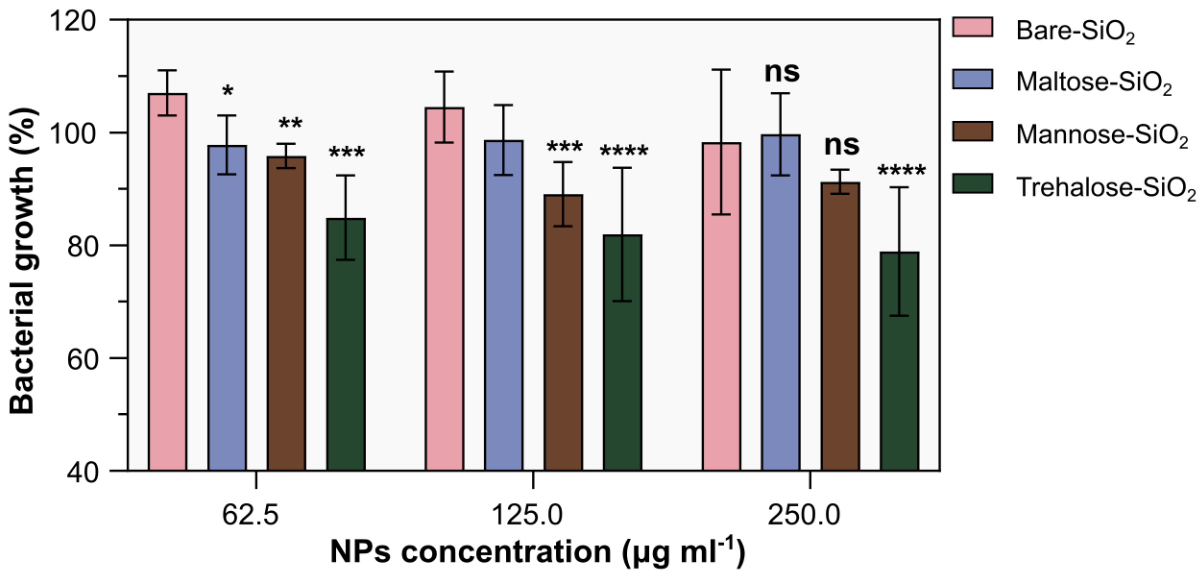


**Figure S6. Bacterial growth following incubation with NPs.** Bacterial growth after 24 h incubation with NPs at concentrations of 62.5, 125, and 250 μg $mL^{-1}$. Statistical analysis was performed using two-way ANOVA to compare carb-$SiO_2$ and bare-$SiO_2$. ns, $P \geq 0.05$; *$P < 0.05$; **$P < 0.01$; ***$P < 0.001$; and ****$P < 0.0001$.

**Table S2. Two-way ANOVA analysis of bacterial growth following incubation with NPs.** Statistical comparison between carb-$SiO_2$ and bare-$SiO_2$ at each nanoparticle concentration.

| | Dunnett's multiple comparisons test | Predicted (ls) mean diff. | 95.00% CI of diff. | Significant? | Summary | Adjusted p value |
|---|---|---|---|---|---|---|
| 62.5 µg ml$^{-1}$ | Bare vs. Mannose | 11.19 | 2.530 to 19.85 | Yes | ** | 0.0075 |
| | Bare vs. Maltose | 9.199 | 0.5406 to 17.86 | Yes | * | 0.0344 |
| | Bare vs. Trehalose | 22.14 | 13.49 to 30.80 | Yes | **** | <0.0001 |
| 125 µg ml$^{-1}$ | Bare vs. Mannose | 15.39 | 6.725 to 24.06 | Yes | *** | 0.0002 |
| | Bare vs. Maltose | 5.825 | -3.107 to 14.76 | No | ns | 0.2836 |
| | Bare vs. Trehalose | 22.55 | 13.88 to 31.21 | Yes | **** | <0.0001 |
| 250 µg ml$^{-1}$ | Bare vs. Mannose | 7.026 | -1.639 to 15.69 | No | ns | 0.1386 |
| | Bare vs. Maltose | -1.39 | -10.32 to 7.541 | No | ns | 0.9669 |
| | Bare vs. Trehalose | 19.38 | 10.72 to 28.05 | Yes | **** | <0.0001 |

Results of the cell viability (Alamar Blue) assay are presented in Fig. S7, and statistical analysis of variance (Two-way ANOVA test) is described in Table S3.

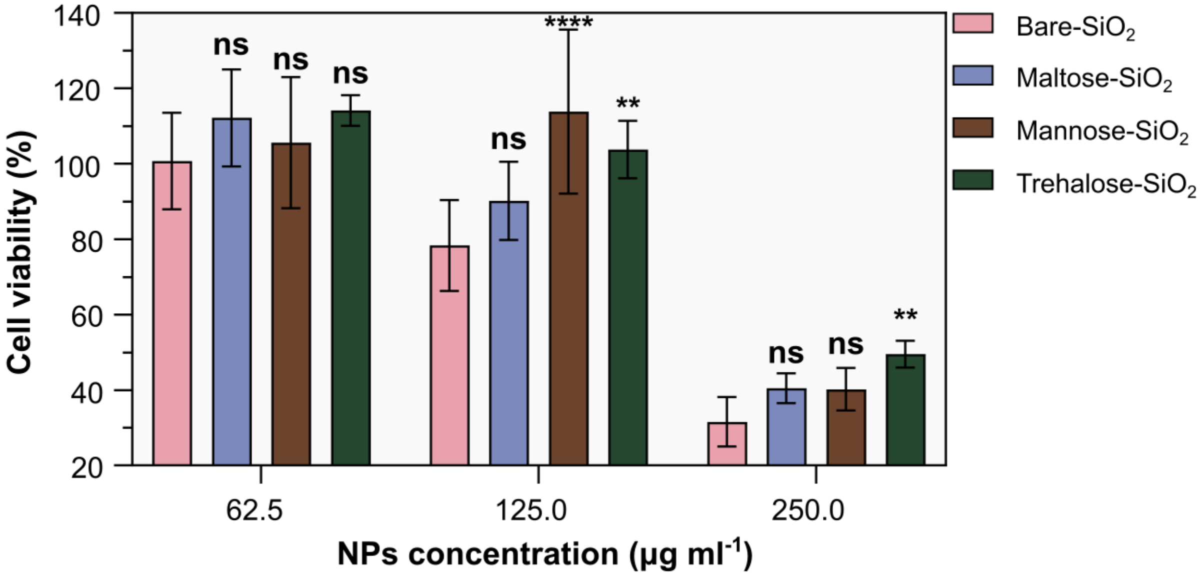


**Figure S7. Cell viability following incubation with NPs.** NIH-3T3 fibroblast viability after 24 h incubation with NPs at concentrations of 62.5, 125, and 250 µg $mL^{-1}$. Statistical analysis was performed using two-way ANOVA to compare carb-$SiO_2$ and bare-$SiO_2$. ns, $P \geq 0.05$; *$P < 0.05$; **$P < 0.01$; ***$P < 0.001$; and ****$P < 0.0001$.

**Table S3. Two-way ANOVA analysis of cell viability following incubation with NPs.** Statistical comparison between carb-$SiO_2$ and bare-$SiO_2$ at each nanoparticle concentration.

| | **Dunnett's multiple comparisons test** | **Predicted (ls) mean diff.** | **95.00% CI of diff.** | **Significant?** | **Summary** | **Adjusted p value** |
|---|---|---|---|---|---|---|
| **62.5 µg ml$^{-1}$** | Bare vs. Mannose | -4.872 | -20.43 to 10.69 | No | ns | 0.7911 |
| | Bare vs. Maltose | -11.44 | -27.00 to 4.121 | No | ns | 0.1945 |
| | Bare vs. Trehalose | -13.45 | -29.01 to 2.106 | No | ns | 0.1042 |
| **125 µg ml$^{-1}$** | Bare vs. Mannose | -35.52 | -52.52 to -18.52 | Yes | **** | <0.0001 |
| | Bare vs. Maltose | -11.84 | -28.11 to 4.437 | No | ns | 0.1998 |
| | Bare vs. Trehalose | -25.45 | -41.73 to -9.174 | Yes | ** | 0.0012 |
| **250 µg ml$^{-1}$** | Bare vs. Mannose | -8.621 | -25.62 to 8.378 | No | ns | 0.4705 |
| | Bare vs. Maltose | -8.9 | -25.18 to 7.376 | No | ns | 0.4112 |
| | Bare vs. Trehalose | -17.91 | -34.18 to -1.631 | Yes | * | 0.0277 |

#### Section S4. Nano-FTIR results

The results obtained from the Imbuia beamline at SIRIUS are presented below. Fig. S8a presents the atomic force microscopy (AFM) image of control *E. coli* (bacteria without NPs incubation), while Fig. S8b displays a representative infrared (IR) spectrum acquired from the region highlighted by the white square in Fig. S8a. The IR spectrum encompasses the three typical regions observed in *E. coli* [14–17]:

I. The protein region, derived from the amide I and II bands, observed between ~1700 and ~1480 $cm^{-1}$;

II. The fatty acid region, ranging from 1480 to 1300 $cm^{-1}$;

III. The bands assigned to $PO_2^-$ stretching at around 1240 and 1080 $cm^{-1}$, predominantly found in nucleic acids.

These biochemical groups are present in the bacterial membrane and DNA. The contents of a Gram-negative bacterial cell are represented in Fig. 4d.

Fig. S8c shows the Gaussian decomposition of the amide I and II bands. The central positions of the Gaussians were determined from maxima of the negative second derivative (window = 5 points, 12.5 $cm^{-1}$ coverage; polynomial order = 3), applied directly to the normalized spectrum without prior smoothing [18] (Fig. S8d). The four colored Gaussian components correspond to the most relevant secondary structure contributions within the amide I band of the *E. coli* control sample, located at: 1682.5 $cm^{-1}$ – antiparallel β-sheet (green Gaussian); 1671.9 $cm^{-1}$ – β-turn (pink Gaussian); 1654.8 $cm^{-1}$ – α-helix (orange Gaussian); and 1634.7 $cm^{-1}$ – parallel β-sheet (blue Gaussian) [17–22].

Figs. S8e and S8f show the normalized negative second derivatives and absorbances of the other regions: region II, corresponding to the lipid area, and region III, corresponding to the nucleic acid area, respectively.

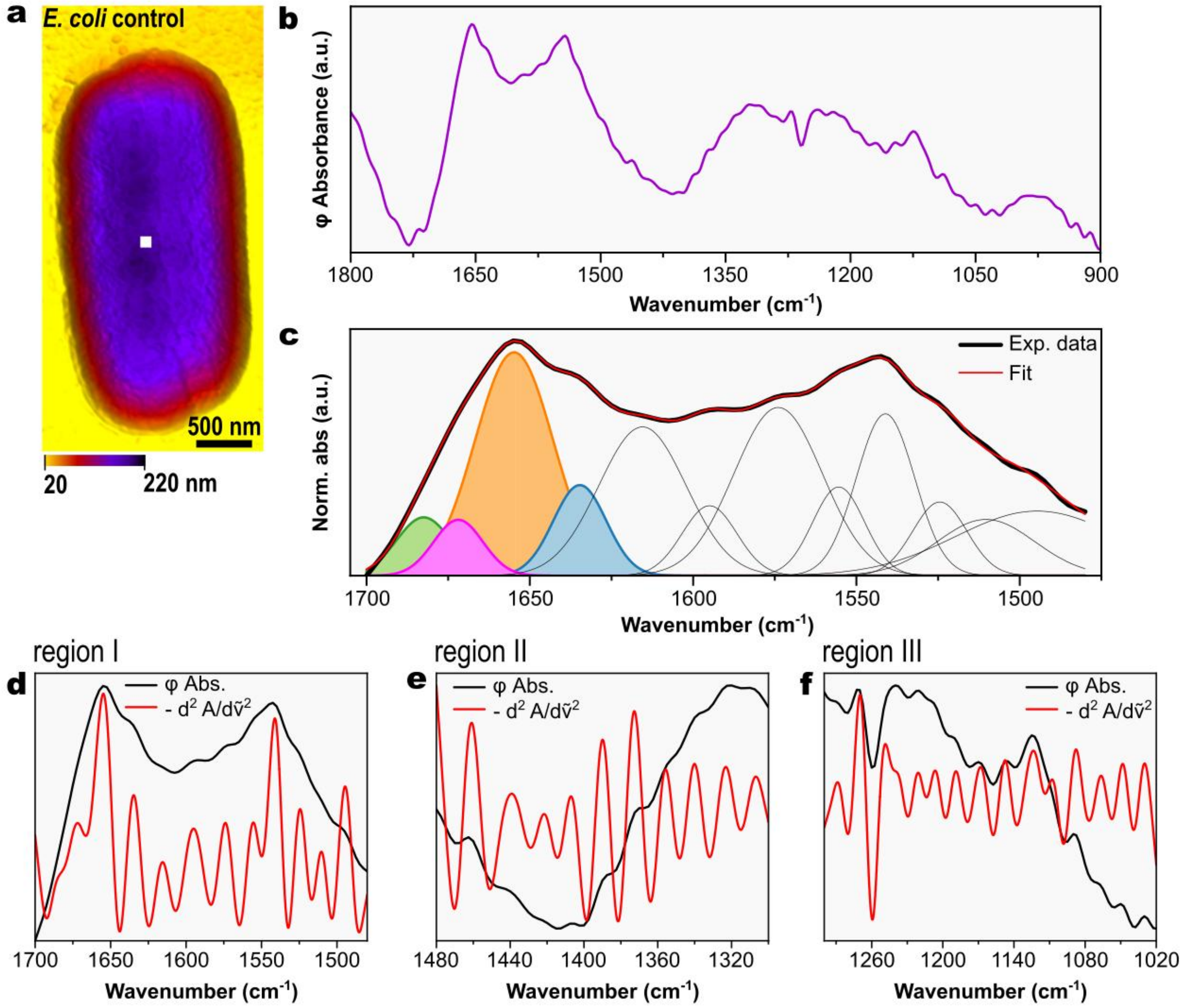


**Figure S8. Nano-FTIR analysis of control *E. coli*.** (a) AFM image of a control bacterium (without NPs). (b) Nano-FTIR spectrum acquired at the location indicated in (a). (c) Deconvolution of the amide I and II bands into Gaussian components centered at the maxima identified from the negative second-derivative spectrum. (d–f) Normalized IR absorbance spectra together with their normalized negative second-derivative spectra for regions I, II, and III, respectively.

An additional spectrum was acquired from the same control sample group to evaluate the reproducibility of the IR measurements, and the resulting spectrum is shown in Fig. S9a.

The deconvolution of the amide I and II bands is shown in Fig. S9b. Section S5 describes the band deconvolution procedure, including the fitting calculations and uncertainty analysis, as well as the parameters used for sample comparison, such as the Iα/Iβ ratio and its propagated uncertainty.

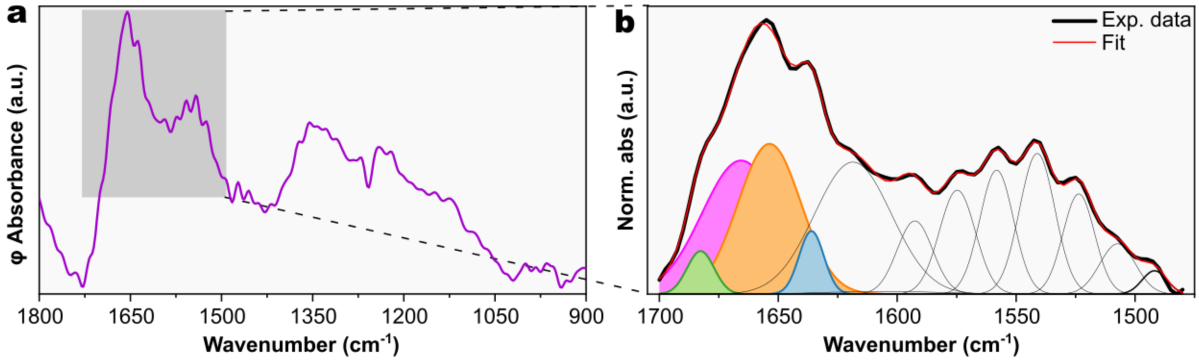


**Figure S9. Additional nano-FTIR analysis of control *E. coli*.** (a) Nano-FTIR spectrum acquired at another location within the control group (without NPs). (b) Deconvolution of the amide I and II bands into Gaussian components centered at the maxima identified from the negative second-derivative spectrum.

The fitting parameters and corresponding Iα/Iβ ratios obtained from the two independently acquired spectra exhibited remarkably similar values, demonstrating a high degree of measurement reproducibility and supporting the robustness of the spectral descriptors used throughout the comparative analysis. This agreement is particularly meaningful in the context of synchrotron infrared nano-spectroscopy (SINS), where spectra are collected from highly localized nanometric regions and each acquisition is intrinsically sensitive to the precise position of the probe. Within this framework, the observed consistency indicates that the spectral signatures identified at the nanoparticle–bacterium interface are not isolated features arising from a single measurement location but instead reflect reproducible interfacial characteristics. Further confidence is provided by the internal coherence of the spectral response across the protein, lipid, and nucleic acid regions, which display mutually consistent trends within each independently acquired spectrum and support the overall interpretation of the molecular changes occurring at the interface.

Figs. S10–S13 display the IR spectra of *E. coli* incubated with bare-$SiO_2$, man-$SiO_2$, mal-$SiO_2$, and tre-$SiO_2$, respectively. Each figure includes the AFM image of the corresponding sample and the IR spectra collected at the highlighted regions in the AFM map: one point above the bacterium, one point above the nanoparticle, and one point at the bacteria–NPs interface.

The amide I and II bands were deconvoluted for spectra acquired on both the bacterial surface and the bacterium–nanoparticle interface to enable direct comparison of the fitted peak positions. The four Gaussian components represent the principal secondary-structure contributions to the amide I band and were assigned consistently across all samples using the fitting strategy established for the control spectrum. The Gaussian components assigned to the α-helix and antiparallel β-sheet structures (orange and blue, respectively) were used to extract the corresponding band intensities ($I\alpha$ and $I\beta$), from which the $I\alpha/I\beta$ ratio was calculated, as presented in the main text (Figs. 5 and 6).

Table S4 summarizes the assigned wavenumbers of the vibrational bands identified in spectra acquired on the bacterial surface for the control sample and each nanoparticle incubation condition.

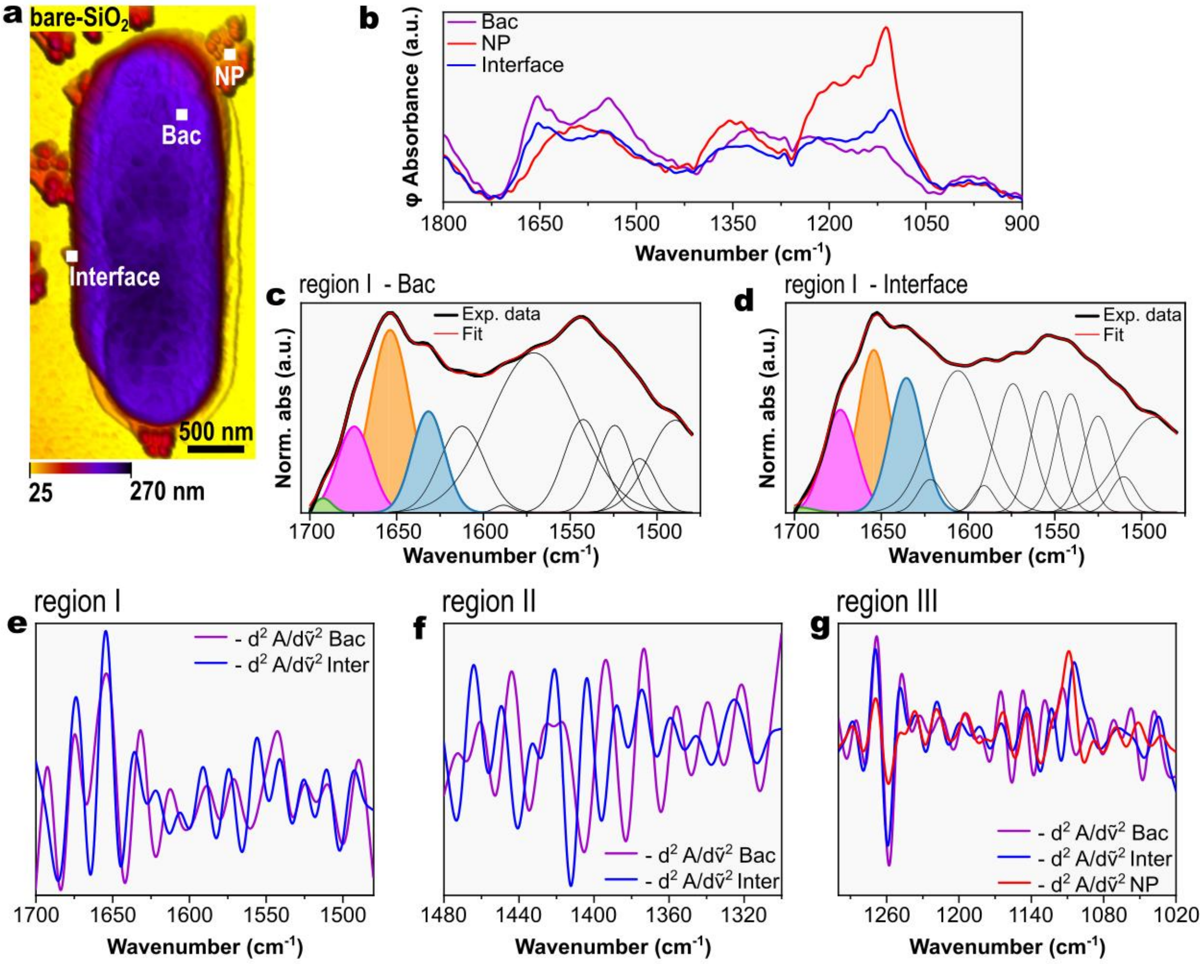


**Figure S10. Nano-FTIR analysis of *E. coli* incubated with bare-$SiO_2$.** (a) AFM image of *E. coli* incubated with bare-$SiO_2$. (b) Nano-FTIR spectra acquired above the bacterial cell, above nanoparticles, and at the bacterium–nanoparticle interface, as indicated in (a). (c) Deconvolution of the amide I and II bands for the spectrum acquired above the bacterial cell into Gaussian components centered at the maxima identified from the negative second-derivative spectrum. (d) Deconvolution of the amide I and II bands for the spectrum acquired at the bacterium–nanoparticle interface into Gaussian components centered at the maxima identified from the negative second-derivative spectrum. (e–g) Negative second-derivative spectra for spectral regions I, II, and III, respectively.

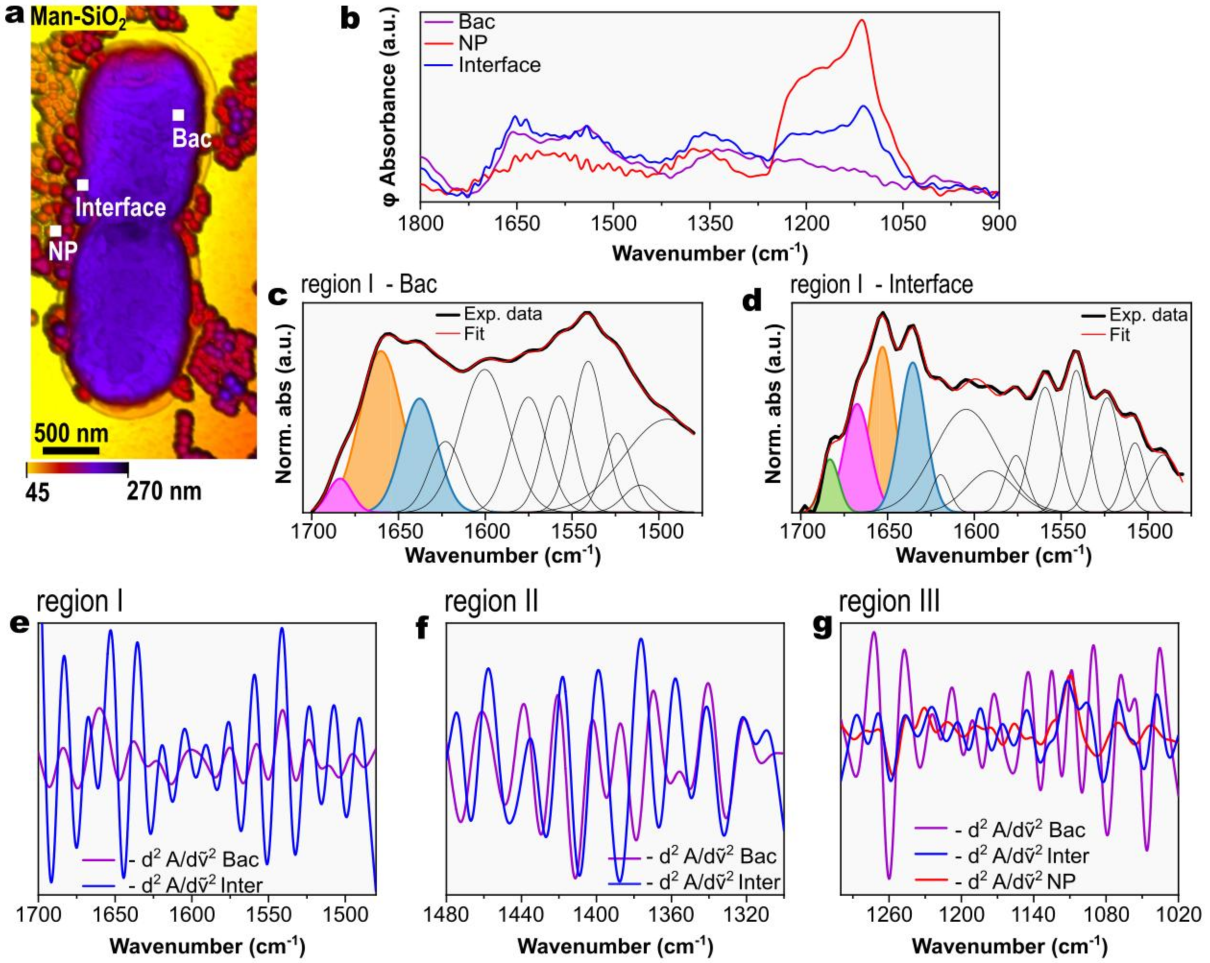


**Figure S11. Nano-FTIR analysis of *E. coli* incubated with man-SiO$_2$.** (a) AFM image of *E. coli* incubated with man-SiO$_2$. (b) Nano-FTIR spectra acquired above the bacterial cell, above nanoparticles, and at the bacterium–nanoparticle interface, as indicated in (a). (c) Deconvolution of the amide I and II bands for the spectrum acquired above the bacterial cell into Gaussian components centered at the maxima identified from the negative second-derivative spectrum. (d) Deconvolution of the amide I and II bands for the spectrum acquired at the bacterium–nanoparticle interface into Gaussian components centered at the maxima identified from the negative second-derivative spectrum. (e–g) Negative second-derivative spectra for spectral regions I, II, and III, respectively.

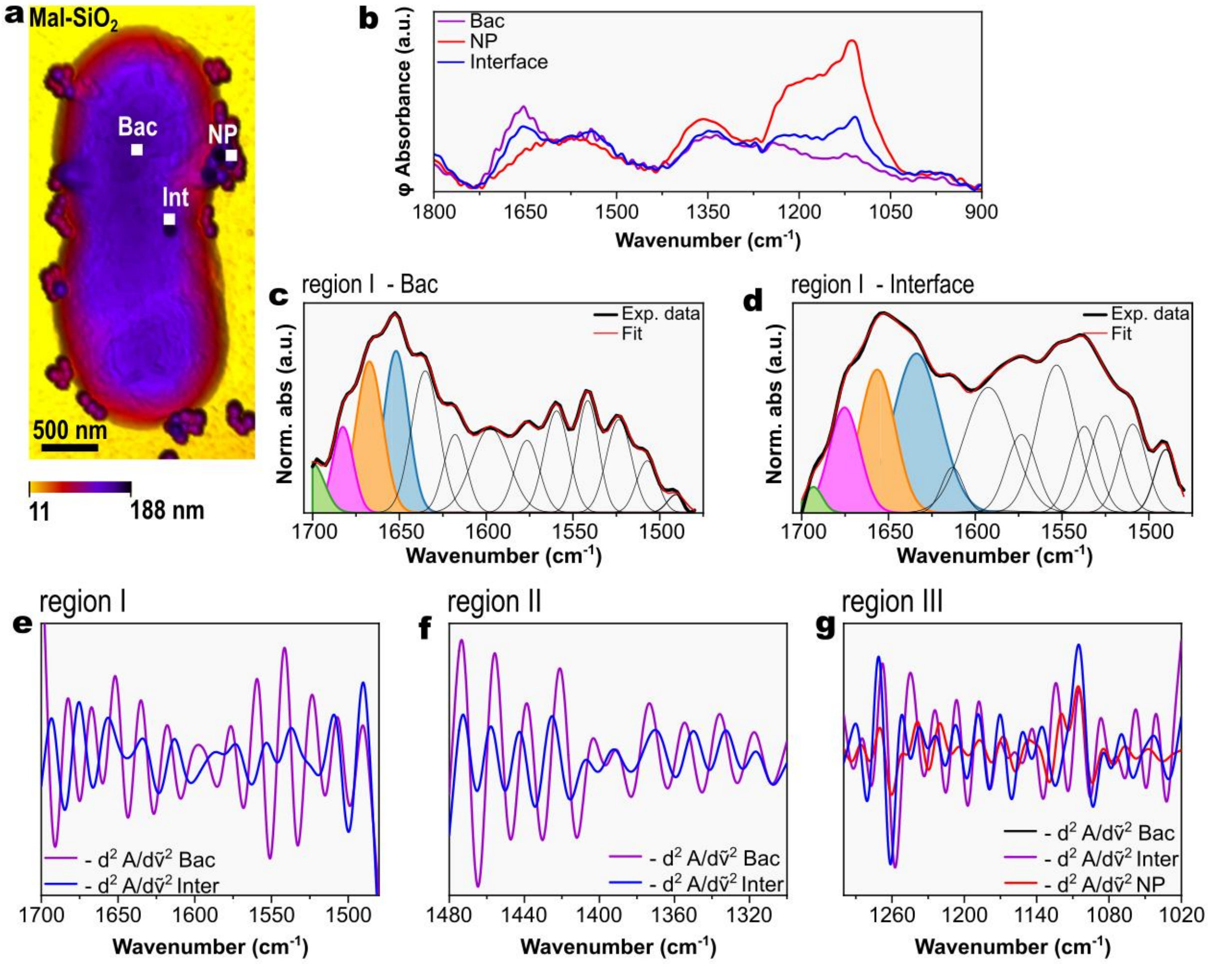


**Figure S12. Nano-FTIR analysis of *E. coli* incubated with mal-$SiO_2$.** (a) AFM image of *E. coli* incubated with mal-$SiO_2$. (b) Nano-FTIR spectra acquired above the bacterial cell, above nanoparticles, and at the bacterium–nanoparticle interface, as indicated in (a). (c) Deconvolution of the amide I and II bands for the spectrum acquired above the bacterial cell into Gaussian components centered at the maxima identified from the negative second-derivative spectrum. (d) Deconvolution of the amide I and II bands for the spectrum acquired at the bacterium–nanoparticle interface into Gaussian components centered at the maxima identified from the negative second-derivative spectrum. (e–g) Negative second-derivative spectra for spectral regions I, II, and III, respectively.

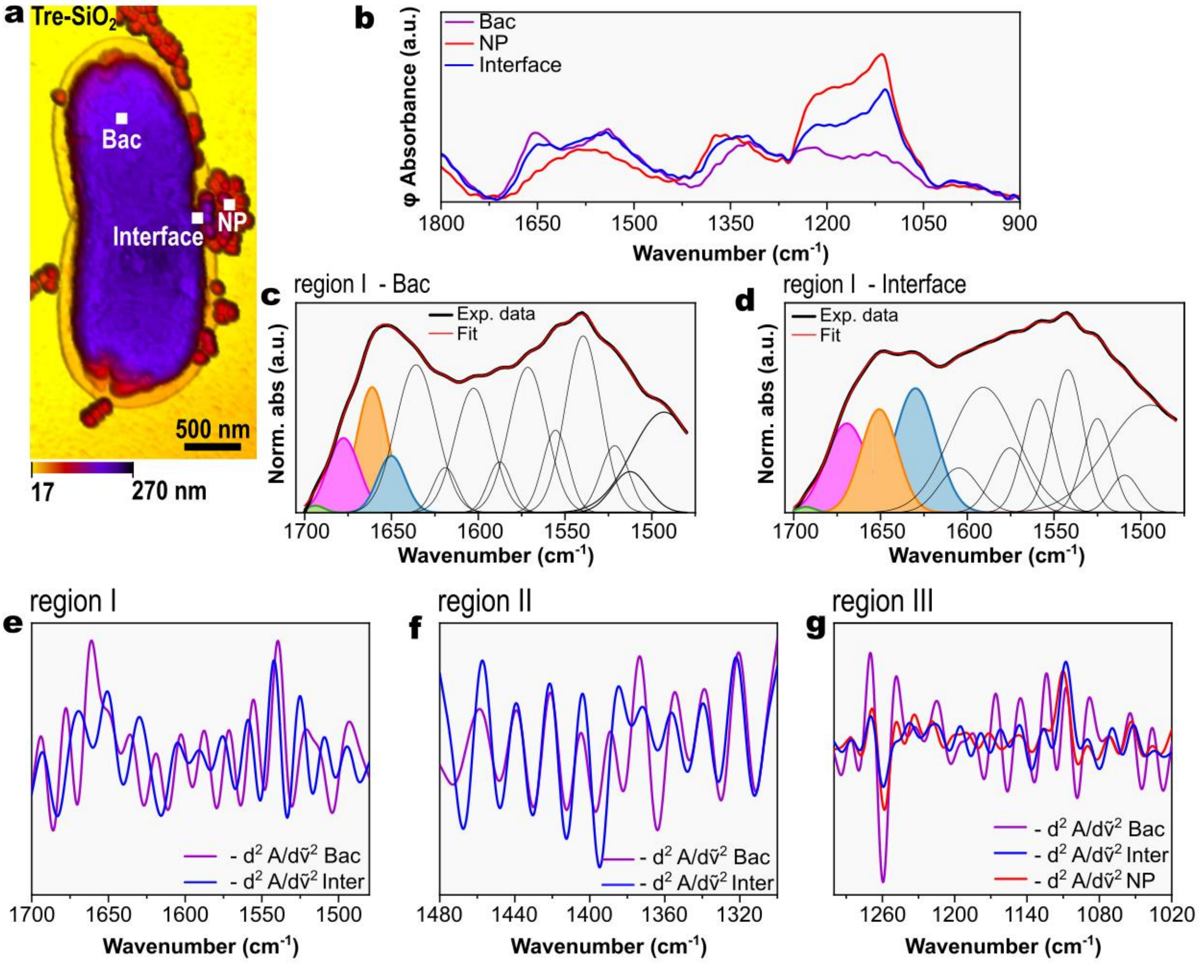


**Figure S13**. **Nano-FTIR analysis of *E. coli* incubated with tre-$SiO_2$.** (a) AFM image of *E. coli* incubated with tre-$SiO_2$. (b) Nano-FTIR spectra acquired above the bacterial cell, above nanoparticles, and at the bacterium–nanoparticle interface, as indicated in (a). (c) Deconvolution of the amide I and II bands for the spectrum acquired above the bacterial cell into Gaussian components centered at the maxima identified from the negative second-derivative spectrum. (d) Deconvolution of the amide I and II bands for the spectrum acquired at the bacterium–nanoparticle interface into Gaussian components centered at the maxima identified from the negative second-derivative spectrum. (e–g) Negative second-derivative spectra for spectral regions I, II, and III, respectively.

Fig. 5f presents the IR spectra collected from regions above *E. coli* cells in the five investigated samples: control bacteria, *E. coli* incubated with bare-$SiO_2$, man-$SiO_2$, mal-$SiO_2$, and tre-$SiO_2$. These spectra are compared to evaluate whether incubation with NPs alone is sufficient to induce alterations in the bacterial membrane structure. Fig. S14 shows the corresponding negative second-derivative spectra for the three highlighted spectral regions.

First, the spectral region between 1700 and 1480 $cm^{-1}$ (Fig. S14a) is dominated by a broad band centered at ~1655 $cm^{-1}$, assigned to amide I, and a second broad band centered at ~1545 $cm^{-1}$, attributed to amide II.[17,19–22]. Second-derivative analysis resolves these broad features into multiple components associated with distinct protein secondary structures. The most prominent contributions correspond to the α-helix and β-sheet components within the amide I and amide II regions, as highlighted in Fig. S14a.

The spectrum of the control *E. coli* closely resembles that of *E. coli* incubated with bare-$SiO_2$, whereas noticeable peak shifts are observed for carb-$SiO_2$. These findings support the interpretation presented in the main text, indicating that carbohydrate coating enhances bacteria–nanoparticle interactions at the bacterial outer membrane.

Additionally, Fig. S14a reveals an extra component at approximately 1650 $cm^{-1}$ in the spectrum of *E. coli* incubated with tre-$SiO_2$. This band is assigned to random coil structures, suggesting a local increase in protein conformational disorder, as discussed in the main text.

The second spectral region, spanning 1480–1300 $cm^{-1}$ (Fig. S14b), provides information on fatty acids in the bacterial outer membrane. This region can be divided into two main contributions: the asymmetric $CH_2$ bending mode at ~1460 $cm^{-1}$ and the symmetric $COO^-$ stretching mode at ~1391 $cm^{-1}$ [14–16]. Incubation with carb-$SiO_2$ leads to noticeable perturbations in this region, suggesting alterations in fatty acid and sugar-related components of the cell wall, despite the spectra being acquired at locations where no nanoparticles were present.

Finally, Fig. S14c shows the negative second-derivative spectra in the 1300–1020 $cm^{-1}$ region. The bands at ~1240 and ~1080 $cm^{-1}$, assigned to the asymmetric and symmetric $PO_2^-$ stretching vibrations of phosphate groups in nucleic acids, respectively, are indicated by gray dashed lines [14–16]. No significant shifts are observed for these bands, suggesting that nanoparticle incubation does not induce detectable changes in phosphate-containing cellular components under the conditions investigated.

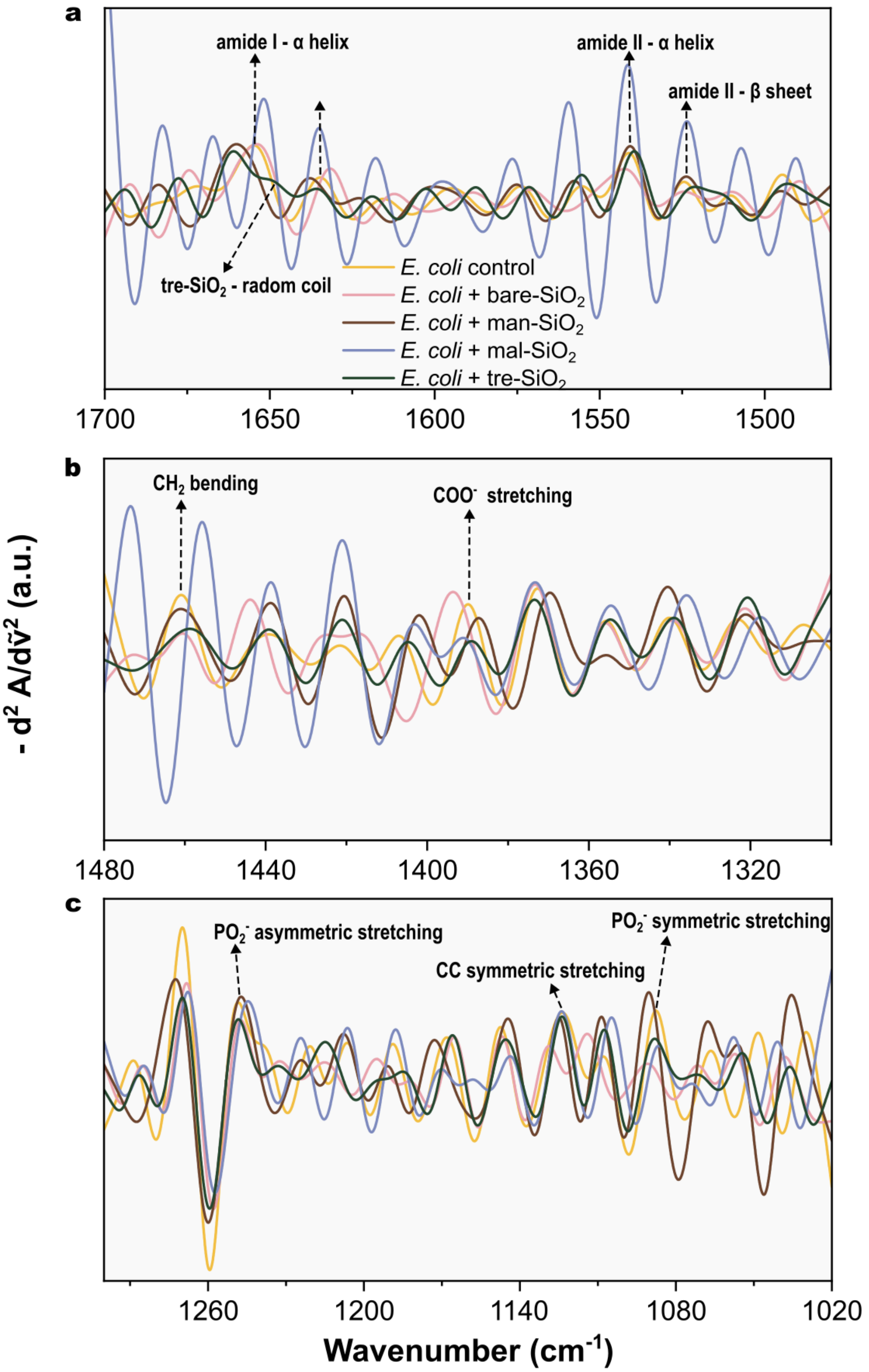


**Figure S14. Negative second-derivative analysis of nano-FTIR spectra acquired above single *E. coli* cells.** Spectra were normalized prior to second-derivative calculation in the following spectral regions: (a) 1700–1500 cm$^{-1}$, (b) 1480–1300 cm$^{-1}$, and (c) 1300–1020 cm$^{-1}$. Yellow, control *E. coli* (without NPs); pink, *E. coli* incubated with bare-$SiO_2$; brown, *E. coli* incubated with man-$SiO_2$; blue, *E. coli* incubated with mal-$SiO_2$; and green, *E. coli* incubated with tre-$SiO_2$.

Table S4 summarizes the vibrational band assignments associated with the main biochemical groups present in the *E. coli* membrane. For each assignment, the corresponding wavenumbers are reported for the control sample and for bacteria incubated with bare- and carb-$SiO_2$, allowing direct comparison of spectral shifts induced by nanoparticle incubation.

**Table S4. Assignment of biochemical groups identified in the *E. coli* membrane.** Biochemical assignments and corresponding wavenumbers for control *E. coli* and *E. coli* incubated with bare-, man-, mal-, and tre-$SiO_2$, based on the spectra shown in Fig. 5f and Fig. S14.

| ***Assignments*** [14–17]* | **Control** | **bare-$SiO_2$** | **man-$SiO_2$** | **mal-$SiO_2$** | **tre-$SiO_2$** |
|---|---|---|---|---|---|
| *Amide I component resulting from antiparallel β -sheets of proteins* | 1682.5 | 1692.4 | 1683.7 | 1682.5 | 1693.8 |
| *Amide I component resulting from β-turn of proteins* | 1671.9 | 1674.5 | - | 1667.2 | 1677.6 |
| *Amide I component resulting from α-helix of proteins* | 1654.8 | 1653.7 | 1660.1 | 1651.9 | 1661.1 |
| *Amide I component resulting from random-coil of proteins* | - | - | - | - | 1650.0 |

| | | | | | |
|---|---|---|---|---|---|
| *Amide I component resulting from parallel β -sheets of proteins* | 1634.7 | 1631.7 | 1637.9 | 1635.1 | 1635.8 |
| *Amide II component resulting from α-helix of proteins* | 1541.1 | 1542.5 | 1540.8 | 1541.5 | 1539.6 |
| *Amide II component resulting from β-sheet of proteins* | 1524.5 | 1524.4 | 1523.8 | 1523.5 | 1521.3 |
| *Asymmetric $CH_2$ bending from lipids* | 1460.9 | 1460.5 | 1461.0 | 1455.7 | 1458.7 |
| *Symmetric $COO^-$ stretching from lipids* | 1389.9 | 1393.6 | 1387.2 | 1391.0 | 1389.1 |
| *$PO_2$ asymmetric stretching from nucleic acids* | 1248.6 | 1247.4 | 1247.3 | 1244.6 | 1248.4 |
| *CC symmetric stretching from ribose* | 1123.6 | 1128.9 | 1125.0 | 1124.0 | 1123.8 |
| *$PO_2$ symmetric stretching from nucleic acids* | 1087.7 | 1090.8 | 1090.4 | 1086.7 | 1088.2 |

#### Section S5. Amide I and II band deconvolution and bootstrap uncertainty estimation

The nano-FTIR spectra in the amide I and II region (1700–1480 $cm^{-1}$) were processed using a three-stage pipeline:

I. **Spectral preprocessing and normalization.** Each spectrum was normalized to its maximum absorbance within the amide I and II region, yielding dimensionless absorbance values in the range [0, 1]. This procedure ensures that the fitted Gaussian amplitudes are directly comparable across samples.

II. **Gaussian band decomposition by nonlinear least-squares fitting.** The center wavenumbers ($\mu_i$) of each Gaussian component were determined prior to fitting through combined analysis of the maxima of the negative second-derivative spectra. These center positions were treated as fixed parameters throughout the fitting procedure. Only the peak intensity ($I_i$, the Gaussian amplitude) and width ($\sigma_i$) of each component were optimized, thereby reducing the dimensionality of the fitting process and minimizing spurious convergence. Here, the Gaussian amplitude is denoted as peak intensity ($I_i$) to distinguish it from the integrated area ($\mathcal{A}_i$).

III. **Statistical uncertainty quantification via residual bootstrap resampling.**

The analysis pipeline was implemented in Python 3 using a custom interactive script based on the SciPy and NumPy libraries. All fitted parameters and associated uncertainty estimates are reported in the Supporting Tables presented in this section.

**Section S5.1 Gaussian decomposition model**

The normalized amide I and II spectrum y(x) is modeled as a superposition of n Gaussian components [23]:

$$y_{fit}(x) = \sum_{i=1}^{n} I_i \cdot exp\left(-\frac{(x-\mu_i)^2}{2\sigma_i^2}\right) \quad \text{(S1)}$$

where x is wavenumber (cm$^{-1}$), $I_i$ is the peak intensity (amplitude) of the i-th Gaussian component, $\mu_i$ is its fixed center (cm$^{-1}$), and $\sigma_i$ is the width constrained to $5.0 \leq \sigma_i \leq 25.0$ cm$^{-1}$. The center positions ($\mu_i$) were fixed using the maxima identified in the negative second-derivative spectra prior to fitting. This approach minimizes parameter correlation associated with strongly overlapping amide bands, stabilizes nonlinear convergence, and improves the reproducibility of comparative spectral analysis across samples [18,24].

The symbol $\mathcal{A}_i$ denotes the integrated area under each component:

$$\mathcal{A}_i = I_i \cdot \sigma_i \cdot \sqrt{2\pi} \quad \text{(S2)}$$

This area is proportional to the relative population of the secondary structure element associated with band i. However, in the present analysis, uncertainty quantification is performed on the peak intensity $I_i$ (Gaussian amplitude), which is the directly fitted parameter and is used to compute the Iα/Iβ ratio reported in Fig. 5g.

The fitting minimizes the sum of squared residuals (SSR):

$$SSR = \sum_{j=1}^{N} \left(y_j - y_{fit,j}\right)^2 \quad \text{(S3)}$$

Minimization was performed using the Trust Region Reflective (TRF) algorithm via `scipy.optimize.curve_fit` (SciPy, Python 3), with up to 20,000 function evaluations. Initial values were set interactively before automated fitting.

Each fit was evaluated by the Root Mean Square Error (RMSE):

$$RMSE = \sqrt{\frac{1}{N}\sum_{j=1}^{N}\left(y_j - y_{fit,j}\right)^2} \quad \text{(S4)}$$

Fits were accepted when the RMSE was commensurate with the baseline noise level. Fits showing systematic residual structure were rejected and re-initialized.

**Section S5.2 Bootstrap uncertainty estimation**

In nonlinear fitting problems, standard errors derived from the linearized covariance matrix can underestimate parameter uncertainties, particularly in the presence of parameter correlations or strong nonlinearity. Bootstrap resampling provides an alternative, model-independent approach that does not rely on local linear approximations and can yield more realistic confidence intervals in such cases [25–27]. In this study, the Gaussian deconvolution of the amide I and II bands involves overlapping components and correlated parameters, making uncertainty estimation nontrivial. Therefore, bootstrap resampling was employed to obtain more reliable estimates of parameter variability.

The residual bootstrap procedure was performed with B = 300 iterations, which limits the meta-statistical error of $\hat{\sigma}$ to approximately 4% [25].

***Step 1 — Compute fit residuals.***

From optimal parameters $\hat{\theta} = \{I_i, \sigma_i\}$:

$$e_j = y_j - y_{fit}(x_j;\, \hat{\theta}), \quad j = 1, \dots, N \quad \text{(S5)}$$

***Step 2 — Generate synthetic spectra by residual resampling.***

For each iteration b = 1, …, B, a perturbation vector e* is drawn with replacement from {$e_1$, …, $e_n$}:

$$y_j^{(b)} = y_{fit}(x_j;\, \hat{\theta}) + e_j^{*} \quad \text{(S6)}$$

This preserves the empirical correlation structure of the residuals without assuming a parametric noise model.

***Step 3 — Re-fit each synthetic spectrum.***

Each synthetic spectrum is re-fitted using $\hat{\theta}$ as the starting point:

$$\theta^{(b)} = arg\, min\theta \sum_{j=1}^{N}\left(y_j^{(b)} - y_{fit}(x_j;\, \theta)\right)^2 \quad \text{(S7)}$$

The optimizer converged successfully in all bootstrap iterations for all analyzed spectra, and no bootstrap samples were excluded from the statistical analysis.

***Step 4 — Compute bootstrap statistics of the peak intensities (amplitudes).***

$$\bar{I}_i = \frac{1}{B}\sum_{b=1}^{B} I_i^{(b)} \quad \text{(S8a)}$$

$$\hat{\sigma}(I_i) = \sqrt{\frac{1}{B-1}\sum_{b=1}^{B}\left(I_i^{(b)} - \bar{I}_i\right)^2} \quad \text{(S8b)}$$

**Section S5.3 Propagation of uncertainty for the Iα/Iβ ratio**

The Iα/Iβ ratio is defined as the ratio between the amplitudes of the α-helical (~1655 $cm^{-1}$) and parallel β-sheet (~1635 $cm^{-1}$) components (Iα and Iβ, respectively) — the two principal protein secondary structures Peak amplitudes were preferred over integrated areas because they are directly optimized during fitting and exhibit lower uncertainty propagation under strong band overlap conditions.

$$R = \frac{\text{I}\alpha}{\text{I}\beta} \quad \text{(S9)}$$

***Bootstrap-Based Uncertainty (Method Used)***

The uncertainty of R was estimated by computing the ratio within each bootstrap iteration:

$$R^{(b)} = \frac{\text{I}\alpha^{(b)}}{\text{I}\beta^{(b)}} \quad \text{(S10)}$$

This implicitly captures the covariance between Iα and Iβ without explicit computation. The bootstrap statistics are:

$$\bar{R} = \frac{1}{B} \sum_{b=1}^{B} R^{(b)} \quad \text{(S11a)}$$

$$\hat{\sigma}(R) = \sqrt{\frac{1}{B-1} \sum_{b=1}^{B} (R^{(b)} - \bar{R})^2} \quad \text{(S11b)}$$

where R̂ is the point estimate from the optimal fit parameters (Eq. S9).

***Comparison with Analytical Error Propagation***

For reference, first-order analytical propagation assuming independence gives:

$$\hat{\sigma}_R{}^{simple} \approx R \cdot \sqrt{\left(\frac{\hat{\sigma}(I\alpha)}{\text{I}\alpha}\right)^2 + \left(\frac{\hat{\sigma}(\text{I}\beta)}{\text{I}\beta}\right)^2} \quad \text{(S12)}$$

The full formula including the covariance term is:

$$\hat{\sigma}_R{}^{full} = \frac{\text{I}\alpha}{\text{I}\beta} \cdot \sqrt{\left(\frac{\sigma(\text{I}\alpha)}{\text{I}\alpha}\right)^2 + \left(\frac{\sigma(\text{I}\beta)}{\text{I}\beta}\right)^2 - \frac{2\, Cov(\text{I}\alpha,\text{I}\beta)}{\text{I}\alpha\cdot\text{I}\beta}} \quad \text{(S13)}$$

where the bootstrap covariance estimator is:

$$\hat{C}ov(\text{I}\alpha, \text{I}\beta) = \frac{1}{B-1} \sum_{b=1}^{B} \left(\text{I}\alpha^{(b)} - \bar{\text{I}}\alpha\right)\left(\text{I}\beta^{(b)} - \bar{\text{I}}\beta\right) \quad \text{(S14)}$$

The bootstrap method (Eq. S10–S11) evaluates the covariance implicitly and is therefore preferred over Eq. S12–S13.

##### Section S5.4 Results

This section reports the RMSE of each fitted band as a measure of fit quality. The peak intensities of the α-helical (~1655 $cm^{-1}$) and parallel β-sheet (~1635 $cm^{-1}$) components (Iα and Iβ, respectively) are also presented, along with their associated uncertainties ($\sigma_\alpha$ and $\sigma_\beta$, reported as absolute values and percentages). Additionally, the Iα/Iβ ratio, its bootstrap mean, and its propagated uncertainty ($\sigma_R$, reported as absolute value and percentage) are provided.

Table S5 presents the results for the control sample, corresponding to the Gaussian fits shown in Figs. S8c and S9b, denoted as Bac 1 and Bac 2, respectively.

Tables S6–S9 present the fit and bootstrap results for *E. coli* incubated with bare-, man-, mal-, and tre-$SiO_2$, respectively, corresponding to the Gaussian fits shown in Figs. S10–S13. Each Supporting Table reports the same parameters described above for two spectral regions: above the bacterial cell and at the bacterium–nanoparticle interface.

**Table S5. Fit and bootstrap results for bacterial control samples.** RMSE values, peak intensities ($I_i$), absolute and percentage uncertainties ($\sigma_i$ and $\sigma_i$%) for the α-helix and parallel β-sheet components, Iα/Iβ ratios, bootstrap means, and ratio uncertainties ($\sigma_R$ and $\sigma_R$%) for Bac 1 and Bac 2, corresponding to the samples shown in Figs. S8c and S9b, respectively.

| **Bac 1 RMSE = 0.005517** | | | | **Bac 2 RMSE = 0.008203** | | | |
|---|---|---|---|---|---|---|---|
| | $I_i$ | $\hat{\sigma}(I_i)$ | $\hat{\sigma}(I_i)\%$ | | $I_i$ | $\hat{\sigma}(I_i)$ | $\hat{\sigma}(I_i)\%$ |
| α-helix | 0.9575 | 0.0105 | 1.1% | α-helix | 0.5512 | 0.0426 | 7.7% |
| β-sheet | 0.3918 | 0.0559 | 14.3% | β-sheet | 0.2394 | 0.0195 | 8.2% |
| Iα/Iβ ratio | | | | Iα/Iβ ratio | | | |
| R | $R_{bootstrap}$ | $\hat{\sigma}(R)$ | $\hat{\sigma}(R)\%$ | R | $R_{bootstrap}$ | $\hat{\sigma}(R)$ | $\hat{\sigma}(R)\%$ |
| 2.4441 | 2.4902 | 0.3484 | 14.3% | 2.3030 | 2.3217 | 0.2468 | 10.7% |

**Table S6. Fit and bootstrap results for bare-$SiO_2$ samples.** RMSE values, peak intensities ($I_i$), absolute and percentage uncertainties ($\sigma_i$ and $\sigma_i$%) for the α-helix and parallel β-sheet components, Iα/Iβ ratios, bootstrap means, and ratio uncertainties ($\sigma_R$ and $\sigma_R$%) for the spectra acquired above the bacterial cell and at the bacterium–nanoparticle interface shown in Fig. S10.

| **bare-$SiO_2$ Bac region RMSE = 0.007056** | | | | **bare-$SiO_2$ Interface region RMSE = 0.007425** | | | |
|---|---|---|---|---|---|---|---|
| | $I_i$ | $\hat{\sigma}(I_i)$ | $\hat{\sigma}(I_i)\%$ | | $I_i$ | $\hat{\sigma}(I_i)$ | $\hat{\sigma}(I_i)\%$ |
| α-helix | 0.9218 | 0.0074 | 0.8% | α-helix | 0.8230 | 0.0141 | 1.7% |
| β-sheet | 0.5126 | 0.0295 | 5.8% | β-sheet | 0.6816 | 0.0303 | 4.4% |
| Iα/Iβ ratio | | | | Iα/Iβ ratio | | | |
| R | $R_{bootstrap}$ | $\hat{\sigma}(R)$ | $\hat{\sigma}(R)\%$ | R | $R_{bootstrap}$ | $\hat{\sigma}(R)$ | $\hat{\sigma}(R)\%$ |
| 1.7984 | 1.8609 | 0.1351 | 7.5% | 1.2074 | 1.2165 | 0.0586 | 4.9% |

**Table S7. Fit and bootstrap results for man-$SiO_2$ samples.** RMSE values, peak intensities ($I_i$), absolute and percentage uncertainties ($\sigma_i$ and $\sigma_i$%) for the α-helix and parallel β-sheet components, Iα/Iβ ratios, bootstrap means, and ratio uncertainties ($\sigma_R$ and $\sigma_R$%) for the spectra acquired above the bacterial cell and at the bacterium–nanoparticle interface shown in Fig. S11.

| **man-$SiO_2$ Bac region RMSE = 0.005059** | | | | **man-$SiO_2$ Interface region RMSE = 0.015988** | | | |
|---|---|---|---|---|---|---|---|
| | $I_i$ | $\hat{\sigma}(I_i)$ | $\hat{\sigma}(I_i)\%$ | | $I_i$ | $\hat{\sigma}(I_i)$ | $\hat{\sigma}(I_i)\%$ |
| α-helix | 0.8133 | 0.0042 | 0.5% | α-helix | 0.8469 | 0.0196 | 2.3% |
| β-sheet | 0.5761 | 0.0107 | 1.9% | β-sheet | 0.7648 | 0.0516 | 6.8% |
| Iα/Iβ ratio | | | | Iα/Iβ ratio | | | |
| R | $R_{bootstrap}$ | $\hat{\sigma}(R)$ | $\hat{\sigma}(R)\%$ | R | $R_{bootstrap}$ | $\hat{\sigma}(R)$ | $\hat{\sigma}(R)\%$ |
| 1.4117 | 1.4170 | 0.0255 | 1.8% | 1.1073 | 1.0790 | 0.0669 | 6.0% |

**Table S8. Fit and bootstrap results for mal-$SiO_2$ samples.** RMSE values, peak intensities ($I_i$), absolute and percentage uncertainties ($\sigma_i$ and $\sigma_i$%) for the α-helix and parallel β-sheet components, Iα/Iβ ratios, bootstrap means, and ratio uncertainties ($\sigma_R$ and $\sigma_R$%) for the spectra acquired above the bacterial cell and at the bacterium–nanoparticle interface shown in Fig. S12.

| **mal-$SiO_2$ Bac region RMSE = 0.008629** | | | | **mal-$SiO_2$ Interface region RMSE = 0.011924** | | | |
|---|---|---|---|---|---|---|---|
| | $I_i$ | $\hat{\sigma}(I_i)$ | $\hat{\sigma}(I_i)\%$ | | $I_i$ | $\hat{\sigma}(I_i)$ | $\hat{\sigma}(I_i)\%$ |
| α-helix | 0.8267 | 0.0177 | 2.1% | α-helix | 0.7230 | 0.0323 | 4.5% |
| β-sheet | 0.7217 | 0.0083 | 1.1% | β-sheet | 0.8013 | 0.0200 | 2.5% |
| Iα/Iβ ratio | | | | Iα/Iβ ratio | | | |
| R | $R_{bootstrap}$ | $\hat{\sigma}(R)$ | $\hat{\sigma}(R)\%$ | R | $R_{bootstrap}$ | $\hat{\sigma}(R)$ | $\hat{\sigma}(R)\%$ |
| 1.1455 | 1.1454 | 0.0318 | 2.8% | 0.9023 | 0.9240 | 0.0493 | 5.5% |

**Table S9. Fit and bootstrap results for tre-$SiO_2$ samples.** RMSE values, peak intensities ($I_i$), absolute and percentage uncertainties ($\sigma_i$ and $\sigma_i$%) for the α-helix and parallel β-sheet components, Iα/Iβ ratios, bootstrap means, and ratio uncertainties ($\sigma_R$ and $\sigma_R$%) for the spectra acquired above the bacterial cell and at the bacterium–nanoparticle interface shown in Fig. S13.

| **tre-$SiO_2$ Bac region RMSE = 0.005523** | | | | **tre-$SiO_2$ Interface region RMSE = 0.005638** | | | |
|---|---|---|---|---|---|---|---|
| | $I_i$ | $\hat{\sigma}(I_i)$ | $\hat{\sigma}(I_i)$% | | $I_i$ | $\hat{\sigma}(I_i)$ | $\hat{\sigma}(I_i)$% |
| α-helix | 0.6362 | 0.0575 | 9.0% | α-helix | 0.5222 | 0.0102 | 2.0% |
| β-sheet | 0.7455 | 0.0351 | 4.7% | β-sheet | 0.6263 | 0.0315 | 5.0% |
| Iα/Iβ ratio | | | | Iα/Iβ ratio | | | |
| R | $R_{bootstrap}$ | $\hat{\sigma}(R)$ | $\hat{\sigma}(R)$% | R | $R_{bootstrap}$ | $\hat{\sigma}(R)$ | $\hat{\sigma}(R)$% |
| 0.8534 | 0.8403 | 0.0817 | 9.6% | 0.8339 | 0.8505 | 0.0454 | 5.4% |

##### Section S5.5 Statistical Comparison of Iα/Iβ Ratios Between Conditions

Statistical comparison of the Iα/Iβ ratios across conditions was performed using a two-tailed z-test based on bootstrap-derived uncertainties. For two measurements with ratio values $x_1$ and $x_2$ and associated bootstrap standard deviations $\sigma_1$ and $\sigma_2$, the test statistic was computed as:

$$z = \frac{|x_1 - x_2|}{\sqrt{\sigma_1^2 + \sigma_2^2}} \quad \text{(S15)}$$

A difference was considered statistically significant at the 95% confidence level when $|z| > 1.96$ (corresponding to $p < 0.05$, two-tailed). It is important to note that $\sigma_1$ and $\sigma_2$ reflect fitting uncertainty estimated by residual bootstrap resampling, therefore the z-test results should therefore be interpreted as a measure of the spectral distinguishability of the fitted ratio values given their fitting uncertainties, rather than as a classical hypothesis test between biological populations.

Table S10 show results for pairwise comparisons of Iα/Iβ ratios. Ratios and standard deviations (σ) are derived from residual bootstrap resampling (B = 300 iterations). Significant differences ($|z| > 1.96$, $p < 0.05$) are indicated with *.

**Table S10. Pairwise z-test comparisons of the Iα/Iβ for *E. coli* incubated with different nanoparticles.** The Iα/Iβ ratios and their standard deviations (σ) were obtained by residual bootstrap resampling. An asterisk (*) indicates a statistically significant difference at the 95% confidence level ($p < 0.05$).

| **Pair** | $x_1$ | $x_2$ | $\sigma_1$ | $\sigma_2$ | **z** | **p-value** | **Significant** |
|---|---|---|---|---|---|---|---|
| **Control *vs* bare-$SiO_2$** | 2.4441 | 1.7984 | 0.3484 | 0.1351 | 1.728 | 0.084 | No |
| **Control *vs* man-$SiO_2$** | 2.4441 | 1.4117 | 0.3484 | 0.0255 | 2.955 | 0.003 | Yes* |
| **Control *vs* mal-$SiO_2$** | 2.4441 | 1.1455 | 0.3484 | 0.0318 | 3.712 | 0.0002 | Yes* |
| **Control *vs* tre-$SiO_2$** | 2.4441 | 0.8534 | 0.3484 | 0.0817 | 4.445 | <0.0001 | Yes* |
| **bare- *vs* man-$SiO_2$** | 1.7984 | 1.4117 | 0.1351 | 0.0255 | 2.813 | 0.005 | Yes* |
| **bare- *vs* mal-$SiO_2$** | 1.7984 | 1.1455 | 0.1351 | 0.0318 | 4.704 | <0.0001 | Yes* |

| | | | | | | | |
|---|---|---|---|---|---|---|---|
| **bare- *vs* tre-$SiO_2$** | 1.7984 | 0.8534 | 0.1351 | 0.0817 | 5.985 | <0.0001 | Yes* |
| **man- *vs* mal-$SiO_2$** | 1.4117 | 1.1455 | 0.0255 | 0.0318 | 6.531 | <0.0001 | Yes* |
| **man- *vs* tre-$SiO_2$** | 1.4117 | 0.8534 | 0.0255 | 0.0817 | 6.523 | <0.0001 | Yes* |
| **mal- *vs* tre-$SiO_2$** | 1.1455 | 0.8534 | 0.0318 | 0.0817 | 3.332 | 0.001 | Yes* |

The only non-significant comparison was between the control and bare-$SiO_2$ groups ($z = 1.728$, $p = 0.084$). All other pairwise comparisons were statistically significant at the 95% confidence level, confirming that the $I\alpha/I\beta$ descriptor is sufficiently sensitive to distinguish the spectral response of *E. coli* to the four nanoparticle surface chemistries investigated. These findings are consistent with those presented in the main text, demonstrating that carbohydrate functionalization of the nanoparticles promotes greater perturbations in the bacterial outer membrane than bare $SiO_2$.

### REFERENCES


(1) Dubois, M.; Gilles, K. A.; Hamilton, J. K.; Rebers, P. A.; Smith, F. Colorimetric Method for Determination of Sugars and Related Substances. *Anal. Chem.* **1956**, *28* (3), 350–356. https://doi.org/10.1021/ac60111a017.

(2) Caracciolo, G.; Palchetti, S.; Colapicchioni, V.; Digiacomo, L.; Pozzi, D.; Capriotti, A. L.; La Barbera, G.; Laganà, A. Stealth Effect of Biomolecular Corona on Nanoparticle Uptake by Immune Cells. *Langmuir* **2015**, *31* (39), 10764–10773. https://doi.org/10.1021/acs.langmuir.5b02158.

(3) Pearson, R. M.; Juettner, V. V.; Hong, S. Biomolecular Corona on Nanoparticles: A Survey of Recent Literature and Its Implications in Targeted Drug Delivery. *Frontiers in Chemistry*. Frontiers Media S. A 2014. https://doi.org/10.3389/fchem.2014.00108.

(4) Galdino, F. E.; Picco, A. S.; Capeletti, L. B.; Bettini, J.; Cardoso, M. B. Inside the Protein Corona: From Binding Parameters to Unstained Hard and Soft Coronas Visualization. *Nano Lett.* **2021**, *21* (19), 8250–8257. https://doi.org/10.1021/acs.nanolett.1c02416.

(5) Ferreira, L. F.; Picco, A. S.; Galdino, F. E.; Albuquerque, L. J. C.; Berret, J. F.; Cardoso, M. B. Nanoparticle-Protein Interaction: Demystifying the Correlation between Protein Corona and Aggregation Phenomena. *ACS Appl. Mater. Interfaces* **2022**, *14* (25), 28559–28569. https://doi.org/10.1021/acsami.2c05362.

(6) Pareek, V.; Bhargava, A.; Bhanot, V.; Gupta, R.; Jain, N.; Panwar, J. Formation and Characterization of Protein Corona Around Nanoparticles: A Review. *J.*

*Nanosci. Nanotechnol.* **2018**, *18* (10), 6653–6670. https://doi.org/10.1166/jnn.2018.15766.

(7) Fleischer, C. C.; Payne, C. K. Nanoparticle-Cell Interactions: Molecular Structure of the Protein Corona and Cellular Outcomes. *Acc. Chem. Res.* **2014**, *47* (8), 2651–2659. https://doi.org/10.1021/ar500190q.

(8) Ahire, J. H.; Chambrier, I.; Mueller, A.; Bao, Y.; Chao, Y. Synthesis of D-Mannose Capped Silicon Nanoparticles and Their Interactions with MCF-7 Human Breast Cancerous Cells. *ACS Appl. Mater. Interfaces* **2013**, *5* (15), 7384–7391. https://doi.org/10.1021/am4017126.

(9) Backus, K. M.; Boshoff, H. I.; Barry, C. S.; Boutureira, O.; Patel, M. K.; D'Hooge, F.; Lee, S. S.; Via, L. E.; Tahlan, K.; Barry, C. E.; Davis, B. G. Uptake of Unnatural Trehalose Analogs as a Reporter for Mycobacterium Tuberculosis. *Nat. Chem. Biol.* **2011**, *7* (4), 228–235. https://doi.org/10.1038/nchembio.539.

(10) Hu, Y.; Liu, X.; Liu, F.; Xie, J.; Zhu, Q.; Tan, S. Trehalose in Biomedical Cryopreservation-Properties, Mechanisms, Delivery Methods, Applications, Benefits, and Problems. *ACS Biomater. Sci. Eng.* **2023**, *9* (3), 1190–1204. https://doi.org/10.1021/acsbiomaterials.2c01225.

(11) Liyanage, S. H.; Yan, M. Maltose-Derivatized Fluorescence Turn-On Imaging Probe for Bacteria Detection. *ACS Infect. Dis.* **2023**, *9* (12), 2560–2571. https://doi.org/10.1021/acsinfecdis.3c00403.

(12) da Cruz Schneid, A.; Albuquerque, L. J. C.; Mondo, G. B.; Ceolin, M.; Picco, A. S.; Cardoso, M. B. Colloidal Stability and Degradability of Silica Nanoparticles

in Biological Fluids: A Review. *Journal of Sol-Gel Science and Technology*. Springer April 1, 2022, pp 41–62. https://doi.org/10.1007/s10971-021-05695-8.

(13) Da Cruz Schneid, A.; Silveira, C. P.; Galdino, F. E.; Ferreira, L. F.; Bouchmella, K.; Cardoso, M. C. Colloidal Stability and Redispersibility of Mesoporous Silica Nanoparticles in Biological Media. *Langmuir* **2020**, *36* (39), 11442–11449. https://doi.org/10.1021/acs.langmuir.0c01571.

(14) Saulou, C.; Jamme, F.; Girbal, L.; Maranges, C.; Fourquaux, I.; Cocaign-Bousquet, M.; Dumas, P.; Mercier-Bonin, M. Synchrotron FTIR Microspectroscopy of Escherichia Coli at Single-Cell Scale under Silver-Induced Stress Conditions. *Anal. Bioanal. Chem.* **2013**, *405* (8), 2685–2697. https://doi.org/10.1007/s00216-013-6725-4.

(15) Hu, X. J.; Liu, Z. X.; Wang, Y. Di; Li, X. N.; Hu, J.; Lü, J. H. Synchrotron FTIR Spectroscopy Reveals Molecular Changes in Escherichia Coli upon Cu2+ Exposure. *Nuclear Science and Techniques* **2016**, *27* (3). https://doi.org/10.1007/s41365-016-0067-9.

(16) Sukprasert, J.; Thumanu, K.; Phung-On, I.; Jirarungsatean, C.; Erickson, L. E.; Tuitemwong, P.; Tuitemwong, K. Synchrotron FTIR Light Reveals Signal Changes of Biofunctionalized Magnetic Nanoparticle Attachment on Salmonella Sp. *J. Nanomater.* **2020**, *2020*. https://doi.org/10.1155/2020/6149713.

(17) Yan, Z.; Li, Q.; Zhang, P. Soy Protein Isolate and Glycerol Hydrogen Bonding Using Two-Dimensional Correlation (2D-COS) Attenuated Total Reflection Fourier Transform Infrared (ATR FT-IR) Spectroscopy. *Appl. Spectrosc.* **2017**, *71* (11), 2437–2445. https://doi.org/10.1177/0003702817710249.

(18) Yang, H.; Yang, S.; Kong, J.; Dong, A.; Yu, S. Obtaining Information about Protein Secondary Structures in Aqueous Solution Using Fourier Transform IR Spectroscopy. *Nat. Protoc.* **2015**, *10* (3), 382–396. https://doi.org/10.1038/nprot.2015.024.

(19) Jackson, M.; Mantsch, H. H. The Use and Misuse of FTIR Spectroscopy in the Determination of Protein Structure. *Crit. Rev. Biochem. Mol. Biol.* **1995**, *30* (2), 95–120. https://doi.org/https://doi.org/10.3109/10409239509085140.

(20) Barth, A. Infrared Spectroscopy of Proteins. *Biochimica et Biophysica Acta (BBA) - Bioenergetics* **2007**, *1767* (9), 1073–1101. https://doi.org/https://doi.org/10.1016/j.bbabio.2007.06.004.

(21) Barth, A.; Zscherp, C. What Vibrations Tell Us about Proteins. *Q. Rev. Biophys.* **2002**, *35* (4), 369–430. https://doi.org/10.1017/s0033583502003815.

(22) Goormaghtigh, E.; Ruysschaert, J. M.; Raussens, V. Evaluation of the Information Content in Infrared Spectra for Protein Secondary Structure Determination. *Biophys. J.* **2006**, *90* (8), 2946–2957. https://doi.org/10.1529/biophysj.105.072017.

(23) Byler, D. M.; Susi, H. Examination of the Secondary Structure of Proteins by Deconvolved FTIR Spectra. *Biopolymers* **1986**, *25*, 469–487. https://doi.org/https://doi.org/10.1002/bip.360250307.

(24) Dong, A.; Huang, P.; Caughey1, W. S. Protein Secondary Structures in Water from Second-Derivative Amide I Infrared Spectra. *Biochemistry* **1990**, *29* (13), 3303–3308. https://doi.org/https://doi.org/10.1021/bi00465a022.

(25) Efron, B.; Tibshirani, R. J. *An Introduction to the Bootstrap*; Chapman and Hall/CRC, 1994. https://doi.org/10.1201/9780429246593.

(26) Kazmierczak, N. P.; Chew, J. A.; Vander Griend, D. A. Bootstrap Methods for Quantifying the Uncertainty of Binding Constants in the Hard Modeling of Spectrophotometric Titration Data. *Anal. Chim. Acta* **2022**, *1227*, 1–10. https://doi.org/10.1016/j.aca.2022.339834.

(27) Sohn, R. A.; Menke, W. Application of Maximum Likelihood and Bootstrap Methods to Nonlinear Curve-fit Problems in Geochemistry. *Geochemistry, Geophysics, Geosystems* **2002**, *3* (7), 1–17. https://doi.org/10.1029/2001gc000253.
